# A multi-scale clutch model for adhesion complex mechanics

**DOI:** 10.1101/2023.01.09.523273

**Authors:** C. Venturini, P. Sáez

**Affiliations:** Laboratori de Càlcul Numèric (LaCaN), Universitat Politècnica de Catalunya, Barcelona, Spain; E.T.S. de Ingeniería de Caminos, Universitat Politècnica de Catalunya, Barcelona, Spain; Institut de Matemàtiques de la UPC-BarcelonaTech (IMTech), Universitat Politècnica de Catalunya, Barcelona, Spain

## Abstract

Cell-matrix adhesion is a central mechanical function to a large number of phenomena in physiology and disease, including morphogenesis, wound healing and tumor cell invasion. Today, how single cells responds to different extracellular cues has been comprehensibly studied. However, how the mechanical behavior of the main individual molecules that form an adhesion complex cooperatively respond to force within the adhesion complex has not been addressed. This is a key aspect in cell adhesion because how these cell adhesion molecules respond to force determines not only cell-matrix behavior but, ultimately, cell function. To answer this question, we develop a multi-scale computational model for adhesion complexes mechanics. Based on the classical clutch hypothesis, we model individual adhesion chains made of a contractile actin network, a talin rod and an integrin molecule that binds at individual adhesion sites on the extracellular matrix. We explore several scenarios of integrins dynamics and analyze the effects of diverse extracellular matrices on the behavior of the adhesion molecules and on the whole adhesion complex. Our results explains how every single component of the adhesion chain mechanically responds to the contractile actomyosin force and show how they control the tractions forces exerted by the cell on the extracellular space. Importantly, our computational results are in agreement with previous experimental data both at the molecular and cell level. Our multi-scale clutch model presents a step forward not only to further understand adhesion complexes mechanics but also to, e.g., engineer better biomimetic materials, repair biological tissues or arrest invasive tumor migration.

**Author summary:** Cell-matrix adhesions are directly implicated in key biological processes such as tissue development, regeneration and tumor cell invasion. These cell functions are determined by how adhesion complexes feel and respond to mechanical forces. Still, how forces are transmitted through the individual cell adhesion molecules that integrate the adhesion complex is poorly understood. To address this issue, we develop a multi-scale clutch model for adhesion complexes where individual adhesion chains, made of integrin and talin molecules, are considered within classical clutch models. This approach provides a rich mechanosensivity insight of how the mechanics of cell adhesion works. It allows to integrate accurate biophysical models of individual adhesion molecules into whole adhesion complex models. Our multi-scale clutch approach allows to extend our current knowledge of adhesion complexes for physiology and disease, e.g., the regeneration of biological tissues or arrest invasive tumor migration, and for engineering better biomimetic materials.

## Introduction

Cell adhesions are central to maintain the right function of the cell [1]. Specifically, integrin-based cell adhesions are responsible for the attachment of cells to the extracellular matrix [2, 3] and are crucial for single and collective cell motility in development and disease [4]. We focus here on integrin-based cell adhesions, and we refer to this type of adhesion complexes (ACs) when we next talk about cell adhesion. ACs are made of an assembling of Cell Adhesion Molecules (CAMs) [5] linked together to establish a rich mechanosensitive and mechanotransductive system that regulates cell adhesion and function [6, 7]. Moreover, ACs grow and respond differently to a number of factors, including the rigidity of the Extracellular Matrix (ECM) [8, 9] and the contractile forces exerted by acto-myosin networks [10]. Today, our knowledge of how ACs respond to these factors is extensive thanks to mounting experimental evidences. In the remaining introductory section, we review the organization of the ACs and their regulation by the ECM.

### Force distribution and architecture in integrin-based cell adhesions

At the nanoscale, ACs are made of ECM ligands, transmembrane and cytoplasmatic proteins and a network of actin filaments and myosin motors (see Fig. 1). The hierarchical composition of the AC was analyzed by measuring the kinematics of each molecule by fluorescence speckle microscopy [11]. Integrins are weakly correlated with the actin flow, indicating that they should be attached to the immobilized ECM. On the other hand, *α*-actinin is fully correlated with the actomyosin network, suggesting that it is fully interconnected within the acto-myosin network. Vinculin and talin show a partial coupling to the actin flow, which indicates that these components connect the acto-myosin structures to the immobile extracellular binding sites. The relation of these components with the actomyosin flow suggested a stratified organization of the main adhesion molecules (Fig. 1). Three-dimensional super-resolution fluorescence microscopy confirmed this organization of CAMs and provided a more precise description of the nanoscale architecture of the AC [5]. They showed that integrins and actin are separated vertically ≈ 40 nm by a first signaling layer consisting of the integrin cytoplasmatic tails, a second layer of talin and vinculin, oriented at 15° with respect to the plasma membrane [12], and an actin regulatory layer containing zyxin and *α*-actinin, among others, that connects talin with the actin network. Therefore, these adhesion molecules lays mostly parallel to the cell-ECM contact plane. The link between integrins, talin, vinculin and the actomyosin network is probably the most determining connection of CAMs for cell-ECM adhesion [7].

**Fig 1.**
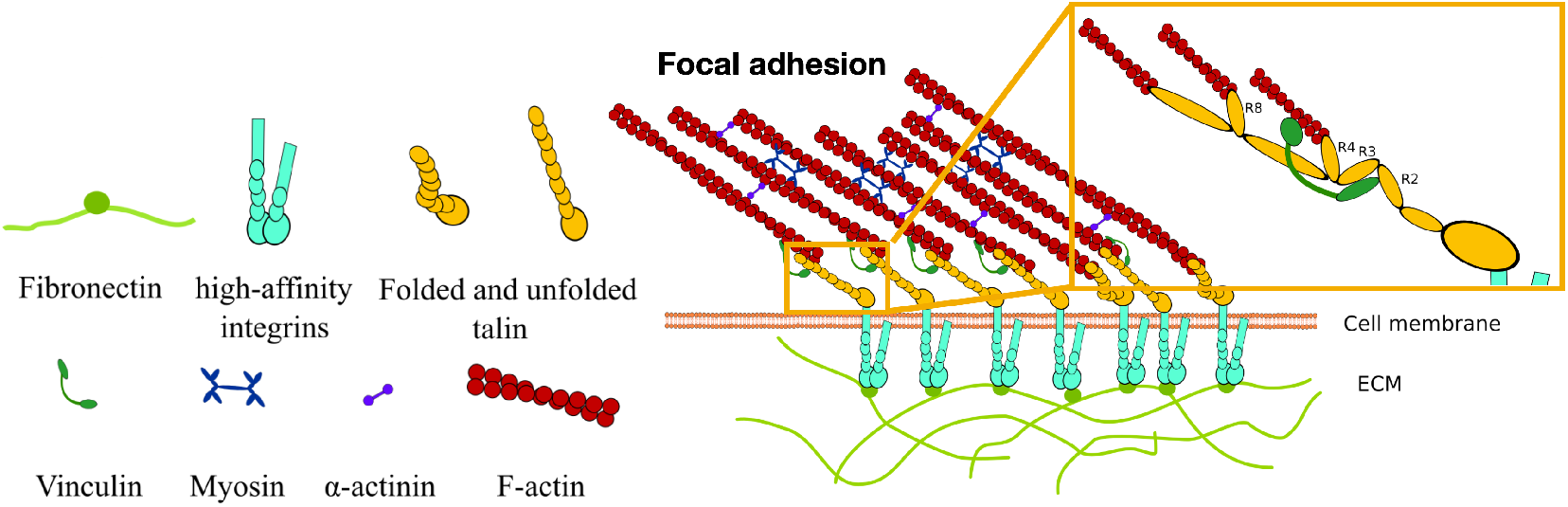
Composition and organization of an AC. On the left, the main cell components of the AC: fibronectin, transmembrane integrins, talin, vinculin and an actomyosin network made of F-actin, myosin motors and *α*-actinin. On the right, the organization of these adhesion complex components. Fibronectin, laminin or collagen serve as anchoring points for the transmembrane integrins to attach to the extracellular space. Integrins bind to the talin rod, which is also connected to the actin network through specific acting binding sites. Actin filaments bundle together mediated by *α*-actinin to form stress-fibers, and exert forces on the adhesion chain by the active pulling forces of myosin motors.

In the inner cellular layer, the *actomyosin network* is made of a template of 10 to 30 actomyosin filaments, linked periodically through *α*-actinin and non-muscle myosin motors that fuse to finally form contractile stress fibers [13, 7, 14, 15]. Myosin motors generate a pulling force that has been implicated in the maturation [16, 7, 16], stability [10, 17] and disassembling [7, 16] of ACs. The force exerted by the myosin motors causes conformational changes in the main CAMs, such as talin and vinculin. Although it is now clear that myosin II plays a key role in the active force generation in cell adhesions [10, 18], how it controls the adhesion dynamics is not completely understood.

The second signaling layer is made of talin and vinculin. *Talin* is recruited from the cytosol to the adhesion site [19] and binds to the intracellular domain of integrins, the *β*-integrin unit, from one side, and to the actin cytoskeleton [20, 21] from its most-inner domain. The force exerted by the actomyosin fibers induces talin deformation and conformational changes [22, 23, 24] (more details on the talin structure and mechanics are discussed in Section). Upon unfolding of the talin rod domains, *vinculin* binds to the exposed talin domains through Vinculin Binding Sites (VBSs) [25]. The binding rates of vinculin to talin increases upon stretching of the talin rod [23, 24].

In the outer molecular layer, transmembrane *integrin* molecules connect the cell to the ECM [26, 3], via laminin, fibronectin or collagen. Integrins diffuse freely within the nascent adhesion. Because the cytoplasmic domains of integrins cannot directly bind to actin, integrins use talin and kidlin as intermediates elements to connect to the actin filaments. Then, upon force-induced activation [19], integrins cluster in the adhesion complex and the diffusivity of the integrins outside the adhesion reduces, which, eventually, leads to the formation of stable ACs [27]. Integrins switch from a low affinity state, where they appear bent and closed, to a high affinity state [28, 29, 30, 31, 32]. Low affinity integrins are activated by their attachment to talin, when they switch to an extended and closed, high affinity state. Integrin activation fosters the binding of integrins with the ECM [33], mediated by talin [34, 35] and vinculin signaling [36]. Once the whole adhesion chain is formed and the force in the chain increases, integrins change again their structure passing to an extended and open state.

Force transmission through all these CAMs is central to the cell adhesion behavior. Force estimations at single integrins have been reported in ≈2-100 pN [30, 3]. Most of these measures were obtained by averaging across the entire adhesion plaque, which only provides an estimation of the actual forces experienced by single integrins. For example, a force value of 1-2 pN at individual integrins was calculated in adhered fibroblasts by estimating the number of bound integrins per unit area [37]. Similarly, a force of ≈10 pN was obtained from traction forces of ≈1 kPa in ≈ 500 integrins / *μ*m^2^ [9]. FRET-based molecular sensors pinpointed the single integrin force in 1-5 pN [38, 39]. DNA-based sensors established a force range of 33-43 pN [29] in *α_v_β*_3_ integrins, one of the most ubiquitous integrin across cell types. A detailed analysis of the lifetime of the *α_v_β*_3_-fibronectin bond showed a maximum lifetime of the bond at 30 pN, vanishing above ≈60 pN [40]. Measures of the force sensed by talin [41] was pinpointed to 7-10 pN and the limit force when all domains are completely unfolded is 30 pN [24].

All these data suggest a narrow mechano-sensing force of few pN across the adhesion chains. Moreover, because the talin-integrin stoichiometry is approximately 1:1 in ACs [5, 42], an integrin and a talin connected in series and stretched from one end share the same force. In this situation, a 30 pN force would unfold all domains of the talin rod [24], and extreme case of the talin state, while the lifetime of the integrin-fibronectin catch bonds would be maximum and, therefore, force could still increase across the molecular chain. Whether this is a realistic, regular scenario in AC behaviour has not been yet shown experimentally.

In summary, there are differences in the force magnitudes measured on these molecules. Whether these differences are real and biologically designed for a right adhesion behavior, which is not consistent with the connection in series of these molecules, or they are discrepancies due to measuring errors, it is not clear.

### The extracellular matrix architecture as a regulator of cell adhesion

The stiffness of the ECM and the spacing of the specific binding sites represent the most determining factors of the ECM for the cell adhesion behavior. The ECM rigidity directly determines the force transmission from the active contraction generated by the acto-myosin network through the CAMs. The behavior of cell adhesion as a function of the substrate rigidity was described by the motor-clutch hypothesis [43, 44, 45, 46]. When integrins bind to rigid substrates, myosin contractility builds force very quickly in each molecular chain and the CAMs must accommodate the imposed displacements. Unbinding rates become faster than binding rates, which limits force transmission to stiff substrates, and the AC completely disengage before other ligands have time to bind. This behavior has been termed ‘‘frictional slippage” [43, 46]. On the other hand, soft substrates deform substantially upon force application, which induce a low force transmission to the CAMs and a cooperative engagement of many bonds along time. As ligands keep binding, myosin contractility deforms the substrate, increasing the force transmission to the substrate, until the bonds reach the limit rupture force. At that point, the AC disassembles, and the cycle starts over. This behavior was previously termed “load-and-fail” dynamics [43, 46].

The AC behavior also depends on the adhesion-site patterning [9, 47]. The ECM is made of proteins, such as fibronectins and laminins, that serve as binding sites for cells to attach. In low ligand spacing and stiff substrates, the maturation of ACs is highly impaired, while in soft substrates ACs consistently mature into long-lived focal adhesions. However, when ligands are put apart by more than few hundreds of nanometers, focal complexes formation is impaired in both soft and stiff substrates [48, 47], probably due to a restriction in integrin clustering, suggesting an adaptor ruler of the order of nanometers that could control cell adhesion maturation. It has also been suggested that the local number and spacing, and not the global density or distribution of ligands, are sufficient to induce adhesion mechanisms [49].

Further research on the cooperative response of ligand spacing and the ECM rigidities in cell adhesion showed contra-intuitive results [50]. On soft substrates, only small nascent adhesions formed for ligand spacing of 50 nm and 100 nm. For these spacing, AC formation was observed only for substrate rigidities above 5 kPa and below 100 kPa, although it was not seen in glass. For 200 nm spacing, AC were observed to form below substrates of 5 kPa. Although most of these results could be described through a ruler mechanism, the formation of ACs for a spacing larger than 200 nm on substrates of ≈ 1 kPa, could not be explained by that hyposthesis.

In short, it is clear that the spacing of ligands and the rigidity of the ECM determine cell behavior. However, how specifically the cooperative mechanical response of the CAMs to these ECM aspects maps to the whole adhesion complex behavior is still not well understood.

### Models for cell adhesion and goal of this work

Different physical models have been proposed in literature to assist current experimental studies for the rationalization of cell adhesion. Continuum models have been used to describe clustering and growth of ACs [51, 52, 53, 54, 55], and the role of the ECM [56] and of cell contractibility [53, 54] in adhesion mechanics. Discrete models have been also used to model cell-ECM mechanics [57, 58, 59]. By corse-grained models, it was also shown that the morphology and distribution of nascent adhesions depend on the nanoscale ligand affinity of *β*-1 and *β*-2 integrins [60]. Other approaches use repeated random sampling simulations, mostly through Monte Carlo (MC) and Gillespie methods, to model the dynamics of cell adhesion [61, 62, 63]. In this framework, the Molecular Clutch model [43, 8, 9] describes the specific binding/unbinding events of individual molecular chains, or clutches, and provides a simple but insightful explanation of cell adhesion mechanics as a function of the substrate rigidity. The clutch models relates the velocity of the retrograde flow of the cell, generated by the pulling forces of myosin motors, with the mechanical properties of the ECM and the clutches dynamics. It was first developed to reproduce the “load and fail” of adhesion in the growth cone of a neuron [43, 9]. The model has been further exploited since then, providing remarkable insights and qualitative understanding of how cell adhesions form, function and modify the whole cell mechanics [8, 64, 65, 66, 67, 68].

Although previous models have explained many experimental observations and have rationalized different aspects of how cell adhesion works mechanically, many fundamental questions remain unsolved. One single AC is an intricate and dynamic compound of molecules, as we summarized above. However, current physical models simplify the actual internal organization of the ACs, which hinders important questions in cell adhesion mechanics. For example, how are the contractile forces transmitted to a single molecular chain? How is the mechanosensing process at individual CAMs during cell adhesion? How do individual molecular chains within the AC behave in different substrate stiffnesses? In this study, we derive a computational multi-scale clutch model to understand how the CAMs that populate an AC behaves during the adhesion process and how they regulate the behavior of the whole AC. In the following sections, we summarize the main features, results and limitations of current clutch models.

### Review of previous molecular clutch models

The clutch model integrates the mechanical response of the substrate and the clutches, that represent the molecular chains, under the contractile pulling of an actomyosin network. Each chain can bind with constant rate and unbind as force builds up on them. This binding and unbinding process is associated with the attachment and disattachment of integrins with the ECM. The solution of the clutch model relies on a repeated random sampling, which is usually solved by Monte Carlo or Gillespie methods. The model computes five main variables involved in cell adhesion behavior [43, 46]: the probability of the clutches of being bound, *P_b_,* the substrate displacement, *x_sub_*, the clutch displacement, *x_c_*, the averaged forces in the clutches, *F_c_*, and the actin network velocity, v. A detailed description of the model is presented in the Methods section.

We first analyze cell adhsion in terms of previous clutch models. We consider ACs crowded with integrins expressing either slip [43] or catch [8] bonds, which we refer to as slip and catch cases from now on. In what follows, and unless specified otherwise, we take model parameters for the on/off rates of slip and catch bonds from literature (see Methods, Table 2). We take values of a slip bond (see Eq. 4) that reproduced experimental data of chick forebrain neurons [43]. To reproduce cells crowded with catch bonds (see Eq. 5), we use model parameters that reproduced the lifetime of *α*_5_*β*_1_ integrins at its maximum activation state in presence of Mn^2+^ ions [40]. The results for the slip and catch cases are in Fig. A1. Our simulations reproduce previous results [43, 8, 46].

**Table 1.**
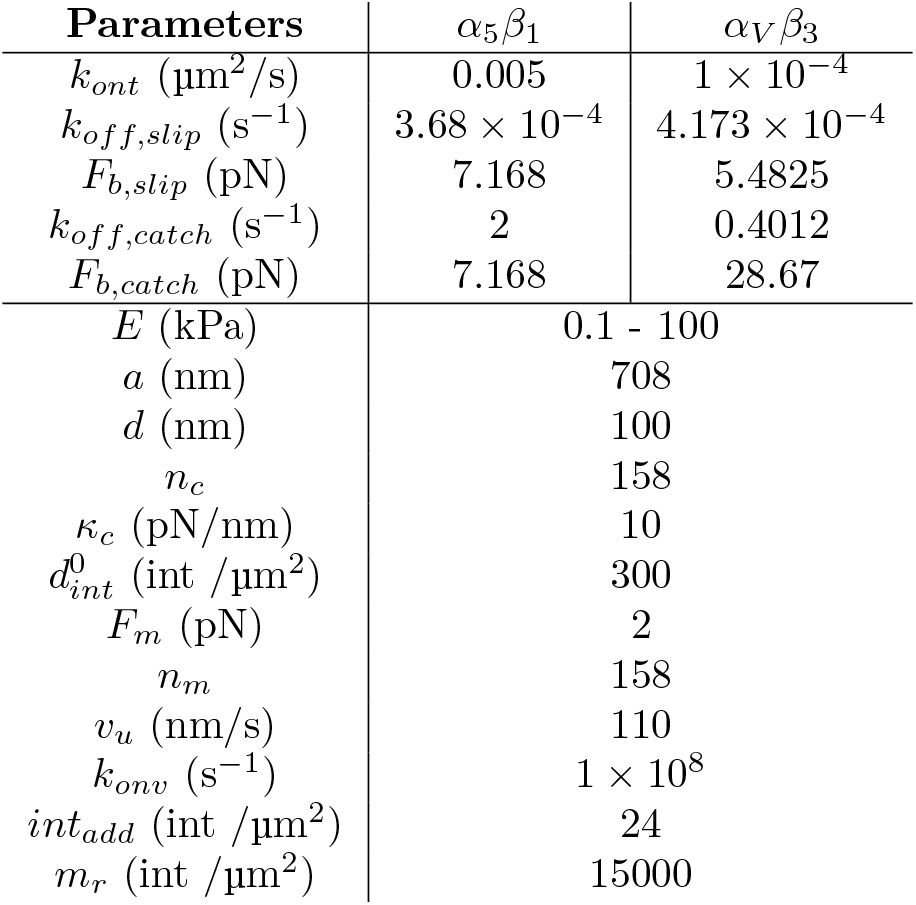
Model parameters for an AC with *α*_5_*β*_1_ and *α_V_β*_3_ integrins.

**Table 2.** Model parameters for slip and catch cases. All model parameters are taken from previous works [43, 8].

| | $E$ | $a$ | $F_{b,slip}$ | $F_m$ | $k_c$ | $k_{off,slip}$ | $k_{ont}$ | $d_{int}$ | $n_c$ | $v_u$ | $n_m$ |
| --- | --- | --- | --- | --- | --- | --- | --- | --- | --- | --- | --- |
| | kPa | nm | pN | pN | pN/nm | s<sup>-1</sup> | $\mu$ m<sup>2</sup>s | intμ m<sup>2</sup> | | nm/s | |
| **Slip**[43] | 0.1 - 100 | 447 | 2 | 2 | 5 | 0.1 | 1 | 1 | 75 | 120 | 75 |
| **Catch** [8] | 0.1 - 100 | 1700 | - | 2 | 1000 | - | 0.0002 | 300 | 1200 | 110 | 800 |

To better understand the model behavior, we analyze the time evolution of all model variables (see Figs. A4 and A5 for details). We specifically focus on the lifetime of each adhesion cycle, that is the average time from the formation of the AC until its complete disengagement. The lifetime of the AC of both slip and catch cases decreases as the stiffness of the substrate increases. In the slip case, the average length of the cycle for *E* = 1 kPa is 7.27 s and for *E* =10 kPa is 0.215 s. In the catch case, for *E* =1 kPa it is 4.25 s and for *E* =10 kPa it is 0.344 s.

We also review recent developments that have extended the original clutch model introducing the effect of talin and vinculin reinforcement [8, 71] (see Fig. A2 for details). Talin unfolding is a mechanosensing event triggered by force that exposes one VBS, where vinculin binds at a force-independent rate *k_onv_*. The unfolding rate of talin, 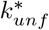, responds to force according to Bell’s model as a slip bond. Talin refolding also depends on force [8]. If talin unfolds, it can either refold again or vinculin can binds to it, increasing integrin density by *int_add_* = 24 integrins/*μm*^2^. The binding rate of integrins follows *k_on_* = *k_ont_d_int_*, where *k_ont_* is the true binding rate characterizing each integrin-fibronectin bond, and *d_int_* is the density of integrins in the AC. If the clutch unbinds before a vinculin binds to talin, integrin density is decreased by *int_add_*, reflecting the fact that adhesions lose integrins if force application is decreased [72, 73]. In the model, integrin density is never allowed to go below the initial value nor above a maximum integrin density, *m_r_* = 15×10^3^ integrins/*μ*m^2^, as there is a limiting density of integrins, or separation between integrins, in the AC (see Fig. A10 for further details). Again, our results reproduce previous experimental and theoretical results [71, 8].

### Limitations in current clutch models for cell adhesion

The molecular clutch model successfully reproduces the adhesion behavior at the cell-scale, specifically the cell traction *P* and the actin velocity *v*, for different cell types (see e.g. [43, 8, 71, 46]). Therefore, it represents an excellent option when we aim at studying the behaviour of an entire cell. However, some parameters and variables of the model contradict previous experimental data. Most probably, this is due to a number of simplifications introduced in the modeling of the adhesion structure.

The first simplification is in the modeling of the substrate. The clutch models consider a single linear spring for the substrate, which naturally results in one single quantity for the substrate displacement. Considering a substrate with nano-patterned attaching locations where integrins adhere to, the displacement of the substrate should change point-wise, depending on the locations of the ligands. This issue has been tackled by including a number of spring elements to represent the ECM [50]. Previous clutch models have simplified the behavior of the chain of CAMs to a linear spring. Integrin and talin, which are key components of the AC [82, 11], are not directly incorporated but have a distinctive mechanical response [24, 3] that contribute significantly to the behavior of the AC. For example, the talin rod undergoes folding and refolding dynamics of all its domains and, moreover, each domain shows a non-linear force-displacement response to load [24].

To better assess the predictions at the molecular scale, we analyzed the deformation of each clutch and compare it to the actual behavior of individual CAMs. Integrins in their bent and closed configuration have a length of ≈ 11 — 13nm [32, 74], called low affinity state. Upon activation, they pass to an extended and open configuration, the high affinity state, with a length of 18 — 23nm [75, 12, 74], from which they can bind and create focal adhesions. *α*_5_*β*_1_ integrins can reach a total length of 50 nm, therefore integrins displacement can reach ≈ 30 nm. Similarly, the talin rod is ≈ 60 —80nm in length [76, 77, 12] in its open, fully folded state and its end-to-end length when fully unfolded under force reaches ≈ 800 nm [24]. This gives a total displacement up to ≈ 730 nm. In the model results, the displacement of the molecular chain has maximum value of ≈ 2 nm (Fig. A2). This displacement is two orders of magnitude smaller than the possible displacements of the integrin and talin molecules.

If the results of the deformations are not well reproduced, one may think that forces at individual molecular clutches could also be poorly reproduced. As we discussed in the introduction, forces have been measured in individual integrins (0 — 40 pN) and talins (0 — 30pN). To analyze the force at each clutch, we plot the mean value and standard deviation as a function of the stiffness of the substrate and the number of ligands *n_c_* for the reinforced case (see Fig. 2). If we neglect the peaks at the end of the cycles, which go up to an unphysical value of ≈ 1500 pN, we see that the forces remain similar for all three values of the substrate rigidity, up to ≈ 60 pN (Fig. 2 and Fig. A2). Although these values are closer to those reported at the individual molecular level, a force of 60 pN would completely unfold all talin domains, which represent an extreme case of the talin behavior.

**Fig 2.**
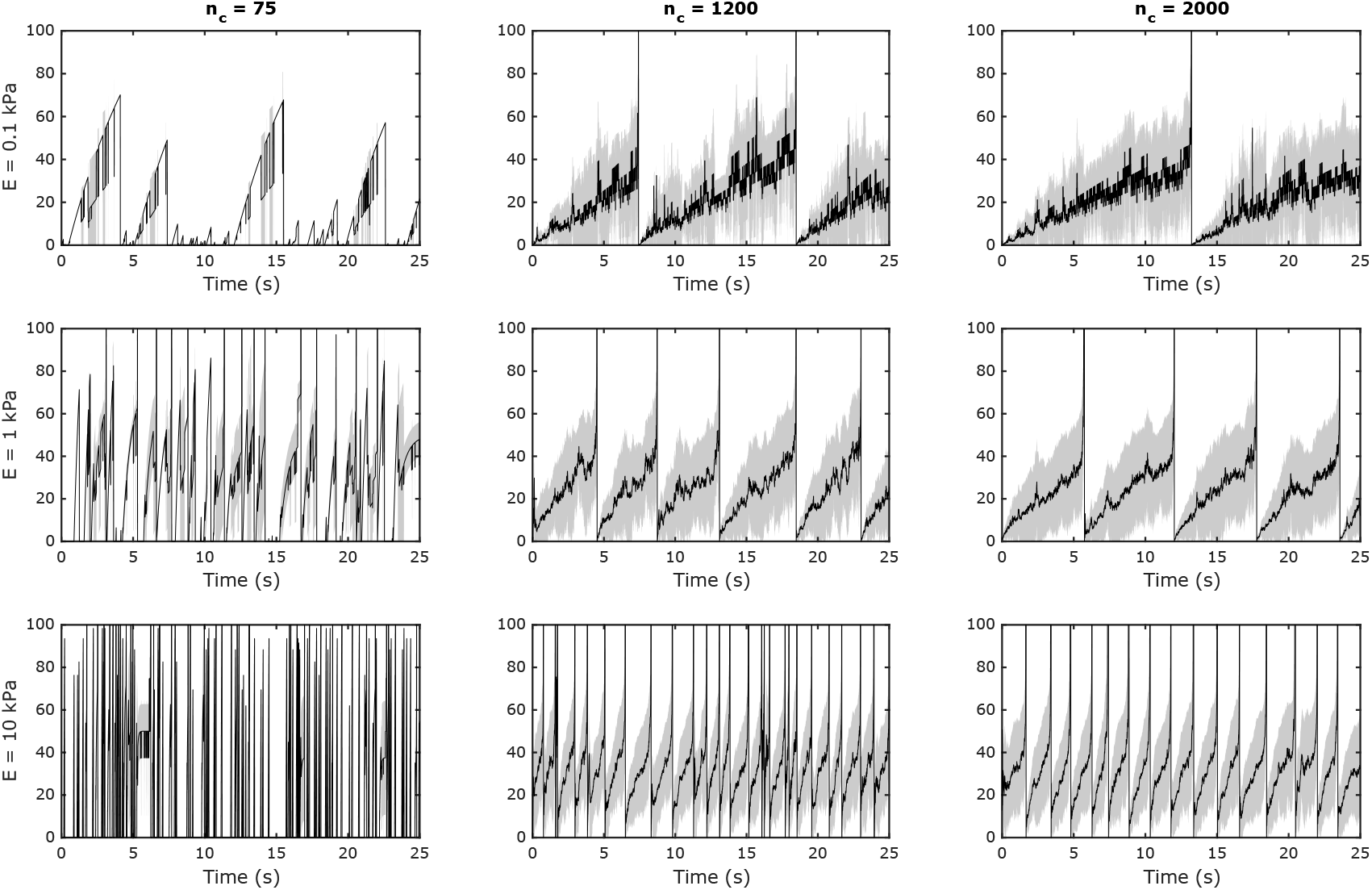
Mean and standard deviation of the force *F_c_* (pN) computed over bound binders. Three values of Young’s modulus (E =0.1, 1 and 10 kPa, in rows) and three values of number of binders *n_c_* = *n_m_* (*n_c_* = 75, *n_c_* = 1200, *n_c_* = 2000, in columns). The rest of parameters are as the default values in Table 2. Standard deviation in light grey. For visualization purposes, the final time is fixed to *t_f_* = 25 s and the y-limit for *F_c_* is set to 100 pN. As the substrate stiffness increases, the average force *F_c_* slightly increases for all three values of *n_c_*. As we increase the substrate stiffness, we see an increase in the number of cycles for a fixed *t_f_* = 25 s. The increase in the averaged force is mostly due to force peaks that appear at the end of each cycle. If the number of cycles increases, as it happens when the stiffness of the substrate increases, the average force increases. Therefore, the peaks obtained at the end of each cycle hide and distort the actual force at the molecular binders.

The results of the lifetime of the AC are also shorter than in previous experimental data. The longest AC lifetime that we obtain is 6.52 s, found for the softest substrate However, previous experimental data show that the time after which the adhesion cluster becomes completely free, the AC lifetime, is ≈ 1 hour [79]. We also observe that the maximum vinculin/talin ratio is 2.5/30 = 0.083. Previous experimental results show vinculin/talin ratios of 1.4 [80] and 2.2 [81]. Perhaps, this is because there are more than one vinculin binding site on the talin rod, which has not been considered in the model.

We hypothesize that an improvement in the molecular characterization within the clutch models could describe more accurately the mechanics of cell adhesion and overcome the limitations and inconsistencies discussed above.

## Results

The main goal of this work is to develop a detailed model of the AC mechanics that may allow us to overcome the limitations discussed above and to further study the mechanosensing mechanisms triggered by the CAMs. To do so, we model a number of molecular chains made of a talin protein attached on top to an actomyosin network and on the bottom to an integrin molecule. We take a previous model of the talin mechanics that considers the conformational changes and non-linear deformation of each domain [24]. Moreover, each integrin molecule links extracellularly to the ECM. Using Green’s functions [83], we compute the deformation of the substrate around each ligand position. Then, we bundle together these individual molecular chains to form an AC (see Fig. 3).

**Fig 3.**
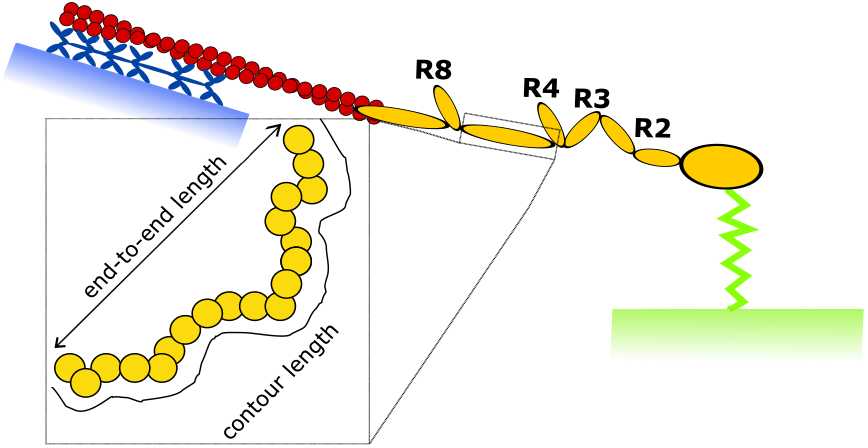
Composition of a single molecular chain. From bottom to top, a single adhesion chain made of a substrate (green), represented as a spring, with one end attached to a fixed surface and the other bound to talin (yellow). The talin’s rod is made of 13 domains and its tail is attached to an actin fibre (red), which is connected to a number of myosin motors (blue), anchored to a fixed surface. Each talin’s domain behaves as a WLC or a FJC in their unfolded and folded states, respectively.

Although we follow the same clutch hypothesis described in previous section, we reinterpret the model. By bundling a number of molecular chains, the new multi-scale clutch model describes the dynamics of a single AC. The macroscopic variables of the model, such as *P* and *v*, represent now the traction under the AC and the velocity of the actomyosin network pulling on the AC. The model is detailed in the Methods section.

### Role of the ECM rigidity on an adhesion chain

To analyze the model behavior and the effect of the talin mechanics in the adhesion behavior, we first model a single adhesion chain where talin attaches to a substrate of various rigidities from one side and remains bound at any time step. We don’t include the integrin molecule between the ECM and talin. On the other end, talin links to an actin filament, that is also bound to *n_m_* myosin motors which are anchored to a fixed surface and generate a contractile force on the actin filament (see Fig. 3).

We run the model through the Gillespie algorithm using as final time *t_f_* = 8 s, which allows for the complete unfolding of the talin domains. We study the same range of substrate rigidities as in previous sections, *E*= 0.1-100 kPa. It is important to note that all model parameters are now based on the individual behavior of each CAM.

We plot the main model variables while varying the Young’s modulus of the substrate (Fig. 4). As we do not allow the chain to disengage, the displacement of the substrate *x_sub_* decreases while increasing the Young’s modulus of the substrate. *x_sub_* is the average over all ligand positions and, therefore, it depends on the displacement at each binder and the total number of bound binders. We do this to obtain substrate displacements comparable to experimental data, which are the average displacement over a certain region of analysis. The resisting force that myosin motors experience also reduces the velocity of the actin filament and, consequently, its total displacement with the increasing rigidity. As the rigidity of the ECM increases, the total displacement imposed in the chain is absorbed by the talin. Therefore, the elongation of talin rod, *x_talin_*, varies from approximately its resting length in the softest substrates (E=0.1 kPa) to ≈350 nm in the stiffest (E=100 kPa). The force on the chain increases from zero to approximately 20 pN in the stiffest substrates. Consequently, the traction in the ECM also increases with the ECM stiffness. Importantly, we show forces and displacements of chain components in agreement with previous data (see again the discussion on the clutch models limitations above).

**Fig 4.**
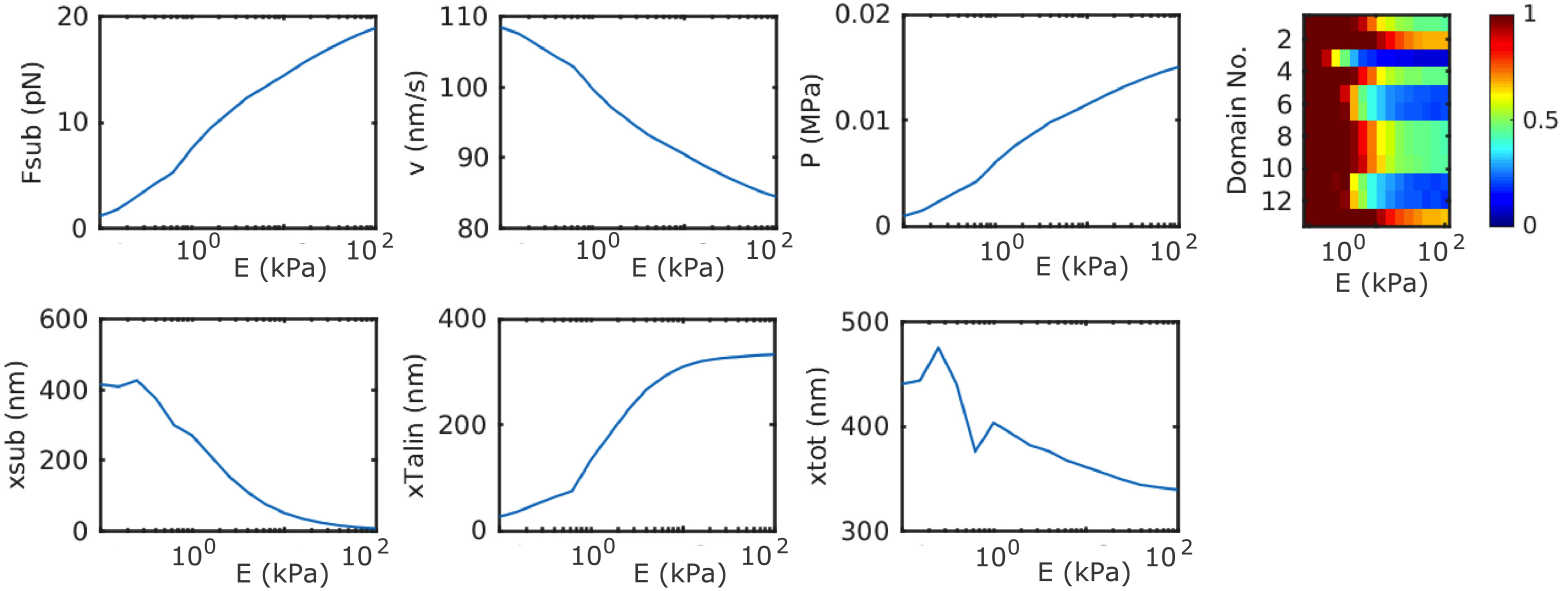
Role of the ECM rigidity on an adhesion chain. Model variables *F_sub_, v, P, x_sub_, x_talin_, x_tot_* for a molecular chain kept bound while varying the Young’s modulus E of the substrate in the range 0.1 — 100 kPa. The color map shows the average folded (red) or unfolded (blue) state for each Young’s modulus and domain of the talin rod.

Talin not only elongates but it also unfolds and refolds during time. At every unfolding event, the length of the talin rod further elongates. We found that, on average, most domains are folded for soft substrates (<10kPa). The unfolding events are mostly triggered starting from a substrate rigidity of ≈ 20 kPa. As we move toward stiff substrates (>30 kPa), the force increases and the domains unfold progressively (Fig. 4), being the R3 domain the most likely one to unfold, followed by the R5, R6, R10 and R11.

### Role of the ECM rigidity in *α*_5_*β*_1_ and *α_v_β*_3_-crowded focal adhesions

Next, we use our multi-scale clutch model to understand how ACs behave. We focus on ACs crowded with either *α*_5_*β*_1_ or *α_v_β*_3_ integrins, among the most ubiquitous types of integrins.

To define the parameters of the integrin dynamics, we look into experimental values for integrin *α_IIb_β*_3_ [88] and assume that *α*_5_*β*_1_ and *α_v_β*_3_ behave similarly in terms of binding rates. Specifically, we take *k_ont_*=0.005 *μ*^2^m s^-1^ and *k_ont_* = 1 × 10^-4^*μ*^2^m s^-1^ for *α*_5_*β*_1_ and *α_v_β*_3_ integrins, respectively. Because *α*_5_*β*_1_ integrins undergoes CMR, we increase its lifetime to reflect the fact that integrins that have been subjected to loading and unloading cycles have a longer lifetime [69]. We do not consider CMR for *α_v_β*_3_ because, as far as we know, it has not been demonstrated for this type of integrins. The lifetime curves together with the experimental data for *α*_5_*β*_1_ [40] and *α_v_β*_3_ integrins [8] are shown in Fig. 5.

**Fig 5.**
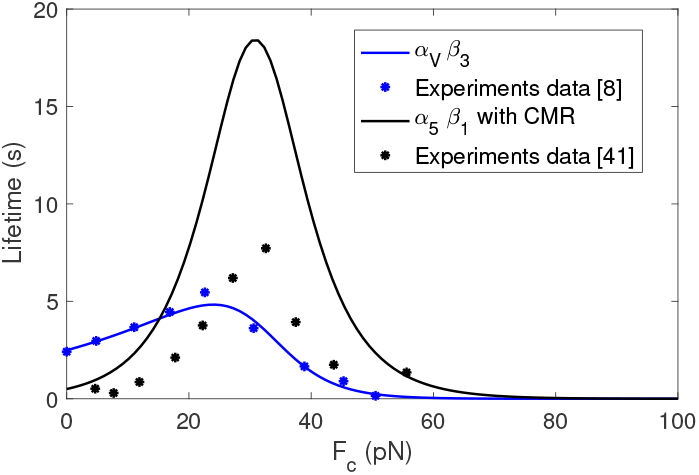
Lifetime for *α*_5_*β*_1_ integrins with CMR and for *α_v_β*_3_ integrins. The parameters obtained for 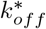 are reported in Table 1

For the Young’s modulus of the substrate, we take again the range *E* = 0.1 - 100 kPa. For the radius of the FA, we take a fixed value of *a* = 708 nm [50] with equispaced ligands at a distance of *d* = 100 nm. The number of equispaced ligands *n_c_* is then computed given the radius of the circular AC and the distance *d*, and we get *n_c_* = 158 binders (see Fig. A10). For the stiffness of the linear spring that models the integrin, we take *κ_c_* = 10pN/nm [40, 69]. For the initial density of integrins, we choose 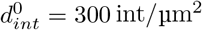 [71, 8]. The stall force of a single myosin motor is *F_m_* = 2pN [89] and we take a number of myosin motors equal to the number of ligands, following previous models [71, 43]. As for the unloaded actin velocity *v_u_*, we choose *v_u_* = 110 nm/s, similar to velocities measured in previous experimental data [90, 91, 92]. We summarize all values for the model parameters in Table 1.

We show the response of the ACs crowded with *α*_5_*β*_1_ and *α_v_β*_3_ integrins as a function of the substrate rigidity in Fig. 6 and Fig. 7, respectively. The cell traction increases with the ECM stiffness, reaching a maximum of ≈ 115 Pa and ≈ 100 Pa for cells crowded with *α_v_β*_3_ and *α*_5_*β*_1_ integrins, respectively, at *E* =10 kPa and remain constant above 10 kPa. The velocity decreases with the increase of the substrate rigidity because of the opposing force to the actin network movement. The minimum velocity, which is again approximately constant above 10 kPa, is ≈ 60 nm/s and ≈ 50 nm/s for the *α_v_β*_3_ and the *α*5*β*1 adhesions, respectively. These traction and velocity values are in agreement with previous clutch models and experimental results [71, 8] (see also previous discussion in Section).

**Fig 6.**
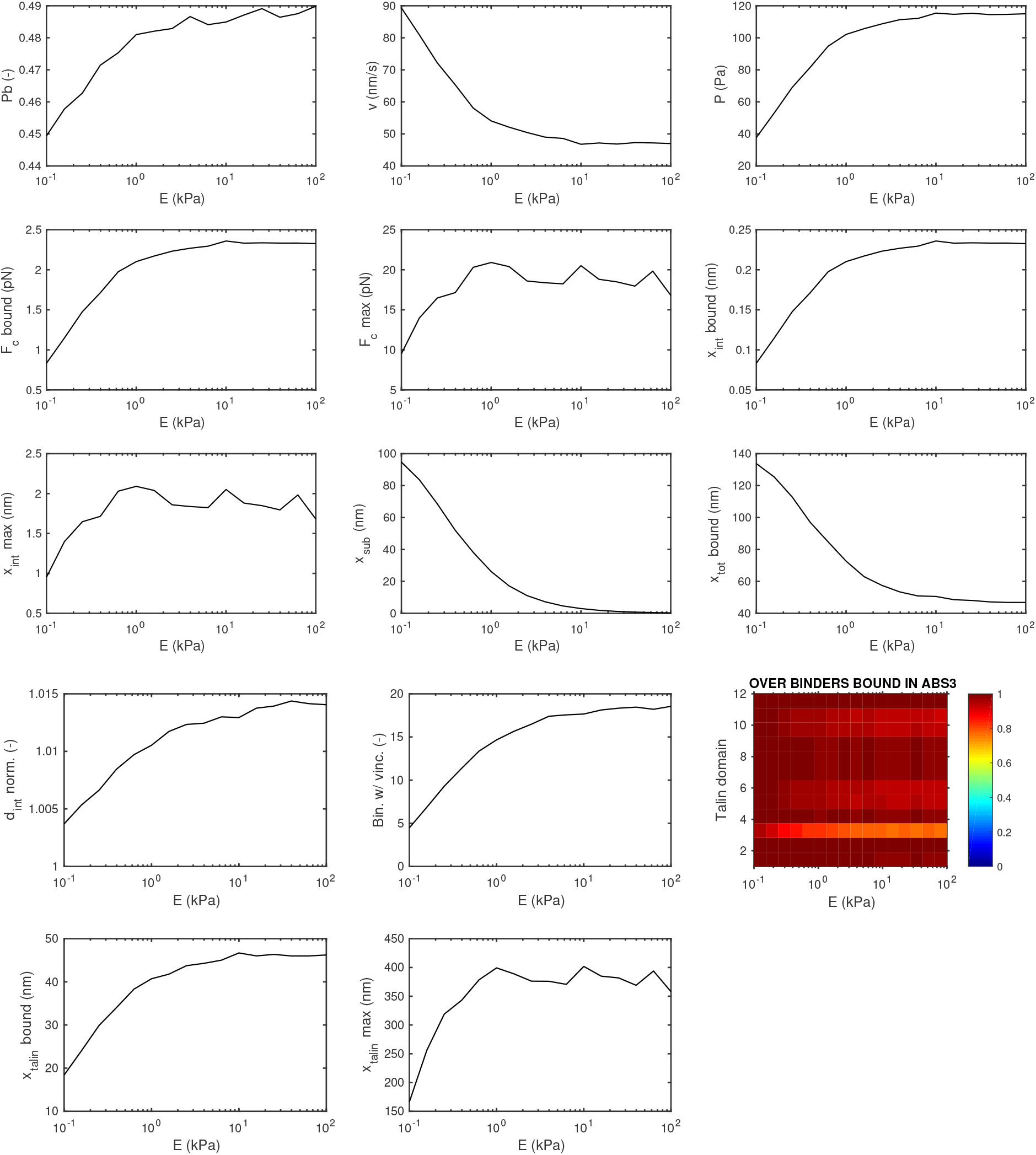
AC with *α*_5_*β*_1_ integrins. Results are averaged over 10 Gillespie simulations. We plot of the main variables against Young’s modulus of the substrate E. The color plot shows the folded/unfolded state of talin domains and it is obtained for one Gillespie simulation, averaging over the bound binders.

**Fig 7.**
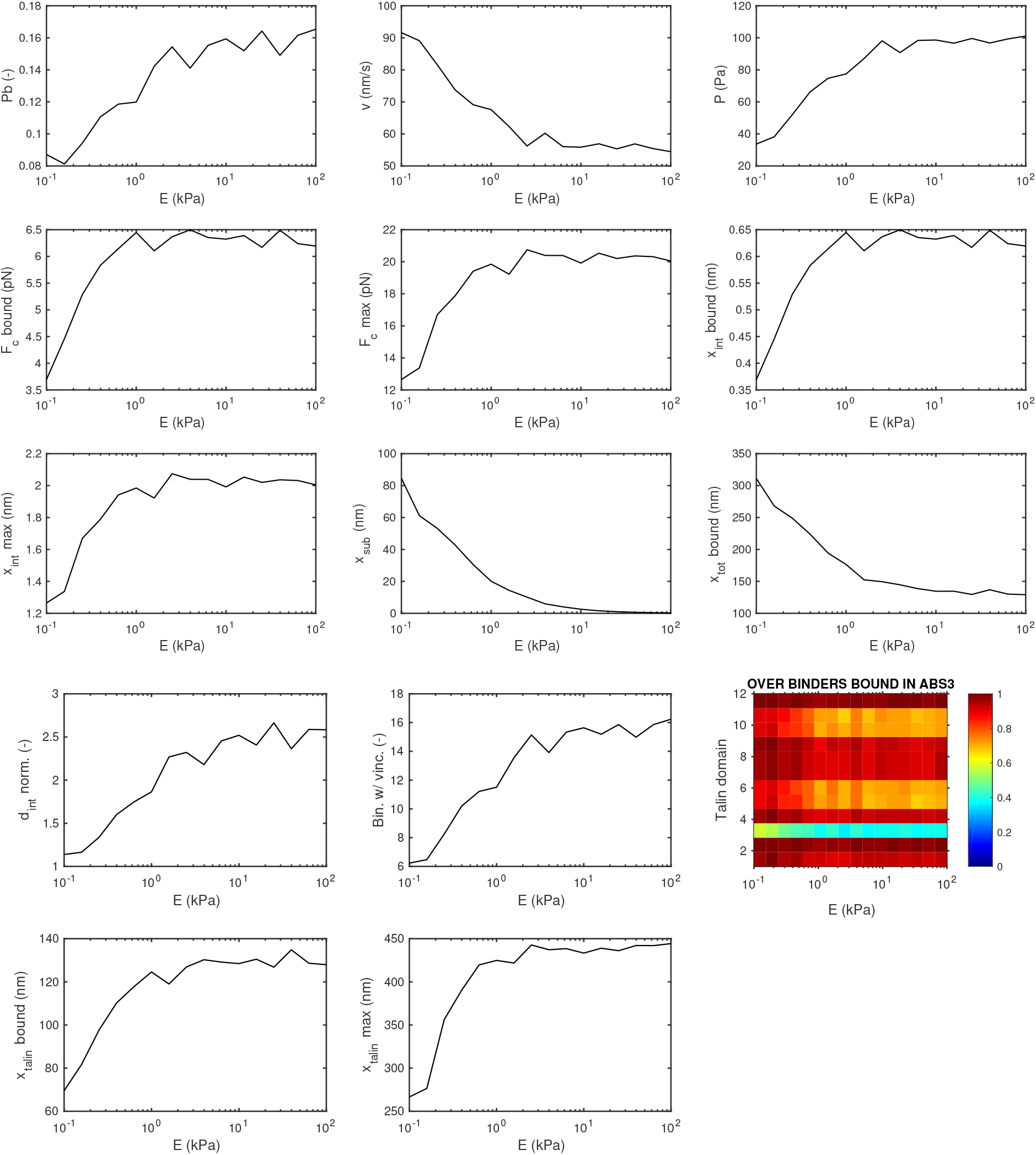
AC with *α_v_β*_3_ integrins. Results are averaged over 10 Gillespie simulations. We plot of the main variables against Young’s modulus of the substrate *E*. The color plot shows the folded/unfolded state of talin domains and it is obtained for one Gillespie simulation, averaging over the bound binders.

The percentage of bound binders increases as the stiffness of the substrate increases, reaching maximum values of 49% and 16% for the *α*_5_*β*_1_ and *α_V_β*_3_ cases, respectively, at E =100 kPa. The lower percentage for the *α_V_β*_3_ case is due to the lower binding rate of this integrin, although its lifetime is larger than *α*_5_*β*_1_ integrins. The maximum force reached at the bound binders, *F_c,max_*, increases from 10-12 pN at 0.1 kPa to 20 pN at 1 kPa in both types of integrins, and remains until 100 kPa. This is in agreement with force values reported at single talin and integrin molecules [24, 40, 69]. The average force over bound binders follows the same increase, with a plateau of ≈ 6.5 pN and ≈ 2pN for the *α_V_β*_3_ and the *α*_5_*β*_1_ cases, respectively, between 10 kPa and 100 kPa. The difference between the two cases is due to the larger lifetime at low forces (<20 pN) and binding rates for *α_V_β*_3_ integrins. The displacement of the ECM decreases as its stiffness increases. The displacement induced by the myosin motors in the molecular chains is mostly absorbed by soft substrates, while integrin and talin molecules absorb the displacement imposed when the substrate increases above the stiffness of these molecules. The maximum displacement of both integrins is ≈ 2*nm*, which is in the order of magnitude of previous experimental results [40, 69]. Again, the average integrin displacement is higher (≈ 0.2nm) for *α_V_β*_3_, aligned with the larger average force. Similarly to other model variables, the displacement of talin increases until substrates of ≈ 10 kPa, and then it remains constant up to the stiffest substrates at ≈ 400 nm and ≈ 440nm for the *α*5*β*1 and the *α_V_β*_3_ cases, respectively. These results are also in close agreement with previous experimental data of the force-displacement relation in the talin rod [24]. Partially, this difference in the talin rod elongation is due to a higher number of unfolded domains in the *α_V_β*_3_ case. Furthermore, the probability of finding a bound binder with a vinculin molecule attached is higher in the *α_V_β*_3_ case because there are more unfolded R3 domains, where vinculin attaches to. Strikingly, the overall number of vinculin molecules in the AC is larger in the *α*5*β*1 case (18) because the number of bound binders is larger. We also see a contraintuitive response in the density of integrins that, as we explained in the introduction, increases according to the increase in vinculin attachments. We show that the integrin density does not change in the *α*5*β*1 while it increases 2.5-fold in the *α_V_β*_3_ case, even though both ACs have a similar increase in vinculin activity. This is because when a binder becomes free, the integrin density increases by *int_add_* if a vinculin is bound to that binder and decreases by *int_add_* if vinculin is not bound. Because *α_v_β*_3_ integrins have a larger lifetime for forces below ≈ 20 pN, most molecular chains do not unbind before the final time *t_f_* and there is not a reduction in integrin density. Therefore, there is not an even correspondence between the value of *d_int_norm* and the total number of vinculins bound for both integrins.

In summary, our results for the tractions and actin flow velocity, which represents the macroscopic variables of the model, are similar to the reinforced case shown in Fig. A2. These are important results because previous clutch models closely reproduce and predict experimental data of these quantities. However, our results also show a close agreement with results of forces and displacements at individual talin and integrin molecules during cell adhesion [24, 40, 69], which represents a clear step forward in the modeling and understanding of the mechanosensing of ACs. It is also worth to note that the two types of integrins analyzed, with two different lifetime landscapes, deliver similar cell tractions. These results indicate that although integrins behavior may change the dynamics of the AC and its mechanosensing response, the resulting cell tractions may not be altered.

### Adhesion dynamics in *α*_5_*β*_1_ and *α_V_β*_3_-based adhesion complexes

To better understand the behavior of ACs crowded with *α*_5_*β*_1_ or *α_V_β*_3_ integrins, we analyze the evolution of a single Gillespie simulation in time. We focus on a substrate rigidity of E = 2.5 kPa (see for results on different substrate rigidities). The model parameters are the same as in previous section.

For the *α*_5_*β*_1_ case, the adhesion enters a quasi-static phase in which the percentage of bound binders stabilizes to ≈ 50% after a short period of ≈ 3 s of transition from the initial free state (Fig. 8). At this quasi-static phase, the actin velocity decreases from the free 110nm/s to ≈ 50 nm/s and the traction at the adhesion patch increases up to ≈120 Pa. All other model variables that are averaged over the bound binders follow the same behavior (Fig. 8). The maximum force reached at a single binders is ≈ 15 pN and presents a peak of 50 pN after which the integrin attachment breaks. The maximum displacement of integrins is ≈ 6 nm. These results are again in agreement with previous data [40, 69]. In terms of the talin dynamics, we show a very rapid landscape of folding and unfolding events which occur every few *ms* (see Fig. 8 and SI). As we showed before, R3 is the domain with more unfolded domains, ≈ 40%, followed by R5, R6, R10 and R11, ≈ 20%. We also see a maximum talin displacement of ≈ 600nm, which represents a value close to talin fully unfolded contour length.

**Fig 8.**
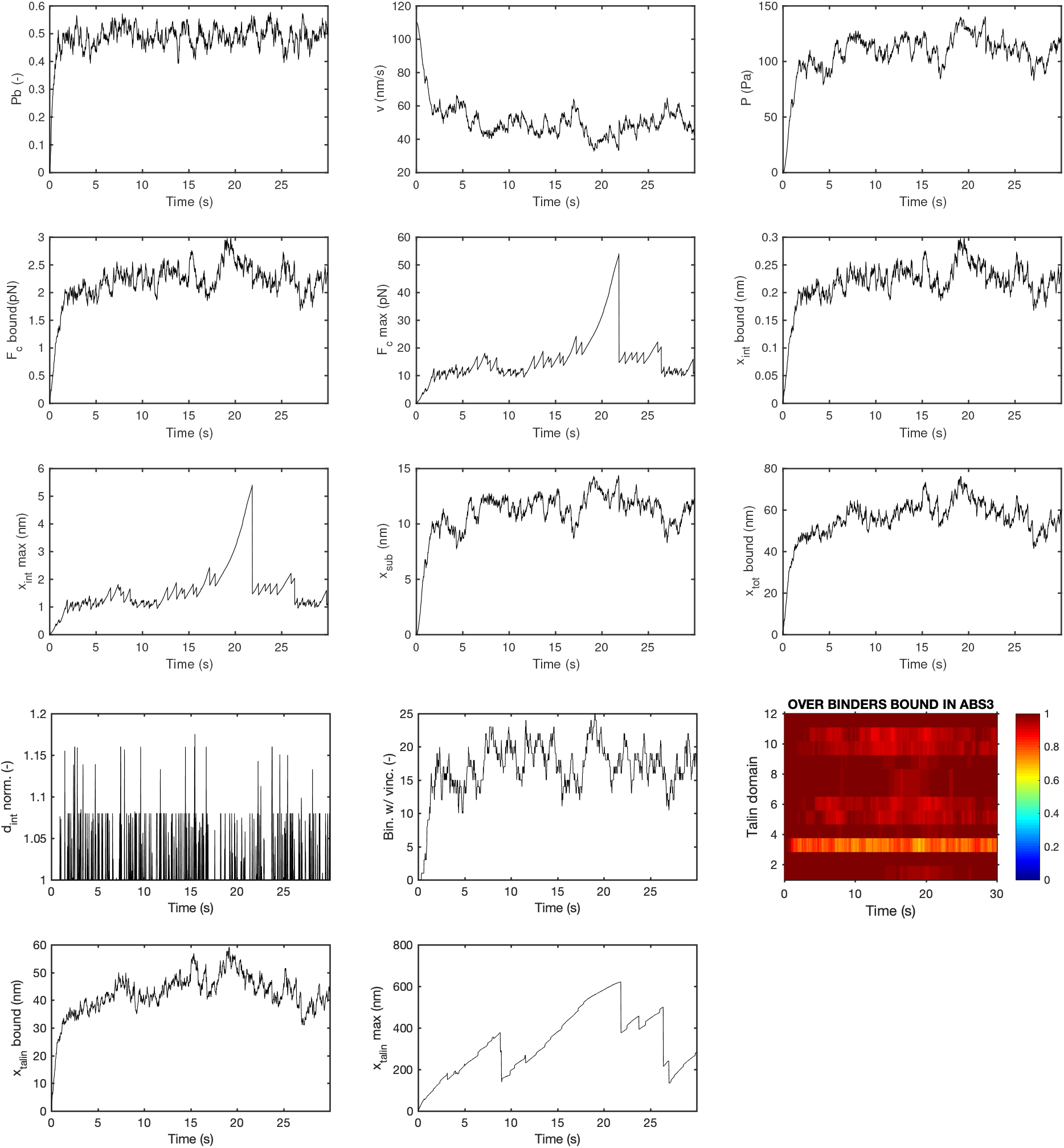
Time behavior of AC with *α*_5_*β*_1_ integrins. Results are for one Gillespie simulation, fixing the Young’s modulus of the substrate to *E* = 2.51 kPa. The figure shows the time evolution for the main variables for the parameters in Table 1.

The results for the *α_v_β*_3_ case have a similar behavior but present some differences. First, the initial transition phase between the free state and the quasi-static state is ≈15s and clutches reach a *P_b_* ≈ 0.16. This is because the larger bonds lifetime cannot compensate the lower binding rates. The velocity and traction at the quasi-static phase reach ≈ 40 nm/s and ≈150 Pa, respectively. The maximum force and displacement present more peaks than in the *α*_5_*β*_1_ case. The peaks reach ≈40 pN and ≈4 nm, respectively, which are lower than in the *α*_5_*β*_1_ case and, again, in agreement with previous data [40, 69]. These differences are due to the larger lifetime of *α_V_β*_3_ integrins below ≈ 20 pN, which explains the amount of peaks around that force value. The talin rod also shows a larger amount of unfolding events in all the talin domains. This is again because the lifetime of *α_v_β*_3_ integrins for forces below ≈ 20 pN is larger than *α*_5_*β*_1_ integrins, which are force values at which most of the talin domains are unfolded. Indeed, we see again that the maximum displacement of talin increases up to ≈ 600 nm, close to the length of the fully unfolded state.

**Fig 9.**
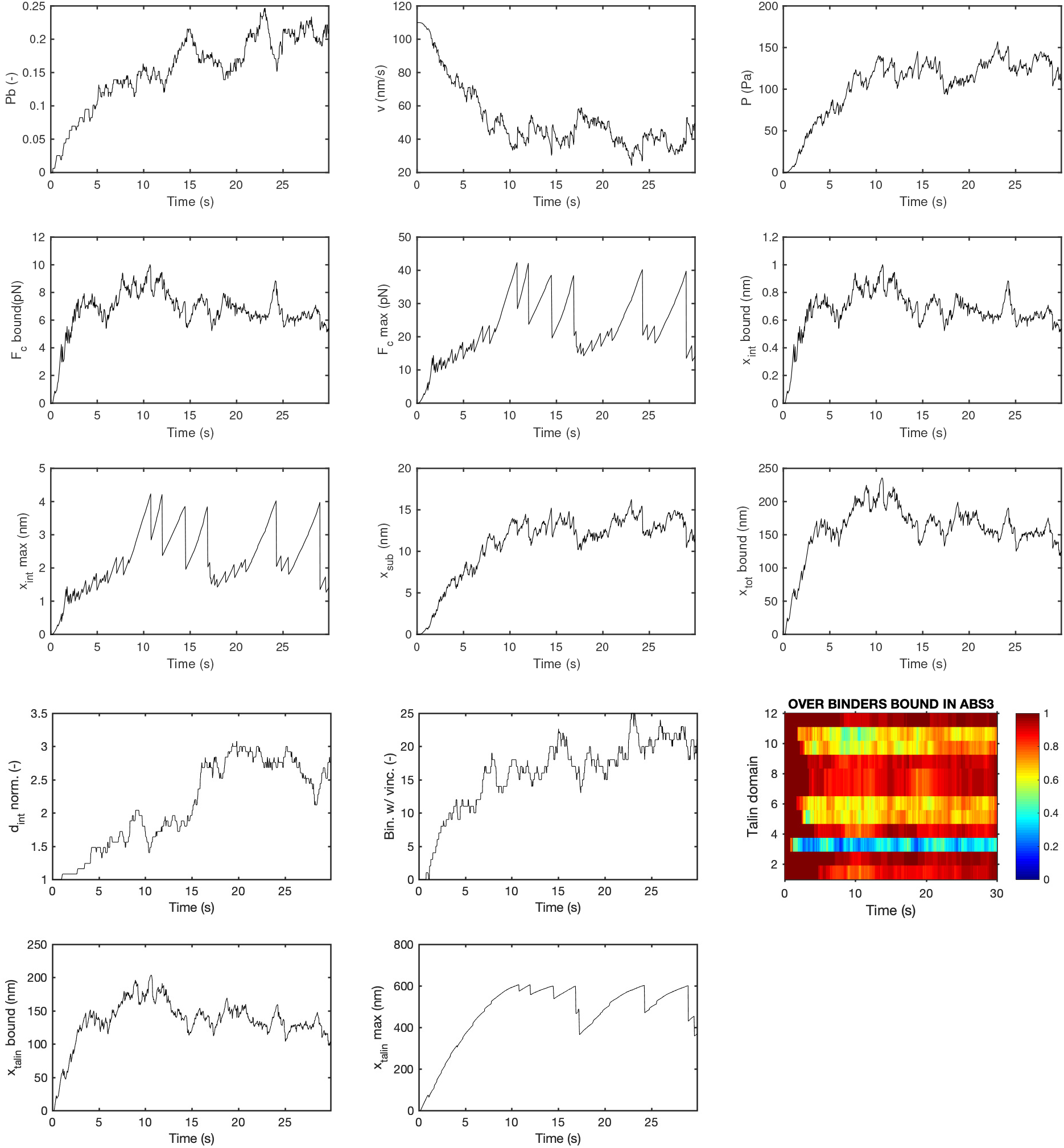
Time behavior of AC with *α_v_β*_3_ integrins. Results are for one Gillespie simulation, fixing the Young’s modulus of the substrate to *E* = 2.51 kPa. The figure shows the time evolution for the main variables for the parameters in Table 1.

As we pointed out above, the mechanosensitivity of both ACs crowded with these integrins gives counterintuitive results. The *α*5*β*1 case has a larger amount of bound binders, which increases the available adhesion chains for vinculin binding. However, there is a limited number of unfolded talin domains. On the other hand, there are also approximately half of the molecular chains available for vinculin binding in the *α_V_β*_3_ case, but there is a larger amount of unfolded domains in the talin rod, that fosters the large number of vinculin attachments. As a result, both cases present a similar increase in bound vinculins. However, as we discussed in the previous section, these increases do not follow a similar increase in integrin recruitment in both cases.

In short, our results show that an AC crowded with *α_V_β*_3_ or *α*5*β*1 behaves very similarly in terms of the traction forces for different substrate stiffnesses. However, our results also suggest that ACs crowded with *α_V_β*_3_ induce a richer mechanosensitivity response compared to *α*5*β*1 integrins in terms of adhesion reinforcement. The unfolding domains of the talin rod and their exposures to VBS further recruits more integrins into the AC, increasing the probability of bound chains. The integrin behavior plays a crucial role in the mechanotransduction of the AC because, when the unbinding rate becomes very fast, there is no time for the increase of force, for the unfolding of talin domains and, therefore, integrin recruitment is severely impaired. These differences may explain the role of different integrins in the mechanosensing of the cell [26].

### Integrins behavior in AC dynamics

Besides *α*_5_*β*_3_ and *α_V_β*_3_ integrins, there are a number of integrins implicated in cell adhesion, e.g. *α*_4_*β*_3_, *α_IIb_β*_3_, *α_V_β*_6_ and *α_V_β*_8_, with a diverse effect in the AC behavior [26]. Even for one specific integrin type, its behavior can change dramatically for different activation states, depending on the presence of different ions [40] or the level of CMR [69]. Consequently, its binding and unbinding rates can be altered. To further explore the effect of different integrin types in the AC behavior, we performed a series of simulations by simultaneously varying the integrin behavior and the substrate rigidity (Fig. 11). The range of parameters related to the integrins behavior and their effects in the lifetime of the bond are shown in Fig. 10. We take the same values of Section for the other model parameters. The effect of increasing the ECM stiffness for all integrins model parameters is an increase in the tractions forces, with the corresponding decrease in the retrograde flow (Fig. 11). Therefore, the effect of the parameters variations in the traction is similar to the previous clutch models and maintains the prediction capability of the clutch model. The increase in cell traction is accompanied by an increase in talin stretch and, therefore, in the number of unfolding events and in the number of vinculin molecules bound to the talin rod. However, as we also discussed above, this does not follow a substantial increase in integrin density.

**Fig 10.**
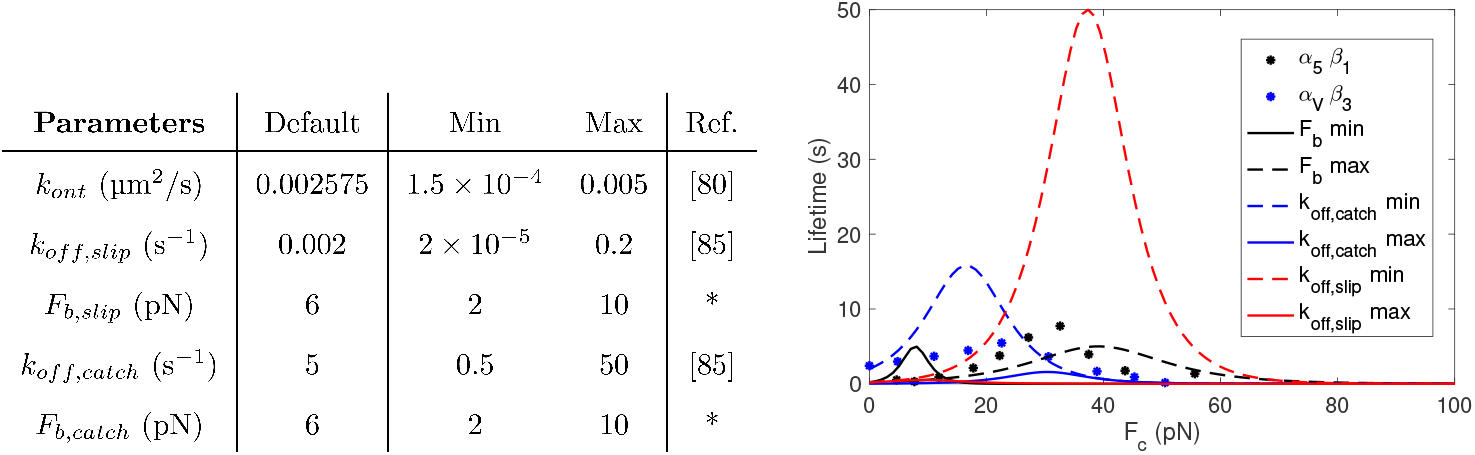
Values of the model parameters. Left: Default values and ranges for the parameters in the binding and unbinding rates of integrins for the multiscale clutch model. * indicates well characterized parameters for which we only analyze a small range of values. For *k_off,slip_* and *k_off,catch_* we adopt an exponential distribution to cover several orders of magnitude. We use a linear distribution for the other parameters. We use previous experimental data on *α_IIb_β*_3_ integrins to define the binding rate [88]. As for the range of the unbinding rate 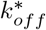, we take into account the experimental data for the lifetime of *α*_5_*β*_1_, *α_V_β*_3_ and *α_L_β*_2_ integrins [93]. Right: Integrin lifetimes obtained with the extremes of the ranges for *F_b,slip_* = *F_b,catch_*, *k_off,catch_* and *k_off,slip_*, together with experimental data for *α*_5_*β*_3_ [40] and *α_V_β*_3_ [8].

We can gather the effects of integrins model parameters in two groups. An increase of *F_b, koff,slip_* or *k_off,catch_* results in a decrease of the bound binders and force transmission, which reduces the cell traction. The decrease in the transmitted force is again followed by a reduction of talin stretch and vinculin attachments. This behaviour is more evident for variations of *k_off,catch_*. These results are expected as we are enhancing the rupture of the bond at low forces by increasing these three parameters. On the other hand, we see an opposite response when we increase *k_ont_*, making the bonds to engage faster than the disengagement. Our results suggest that the on/off rates, which are indicative of different integrins and different activation states, results in very different AC behaviors.

**Fig 11.**
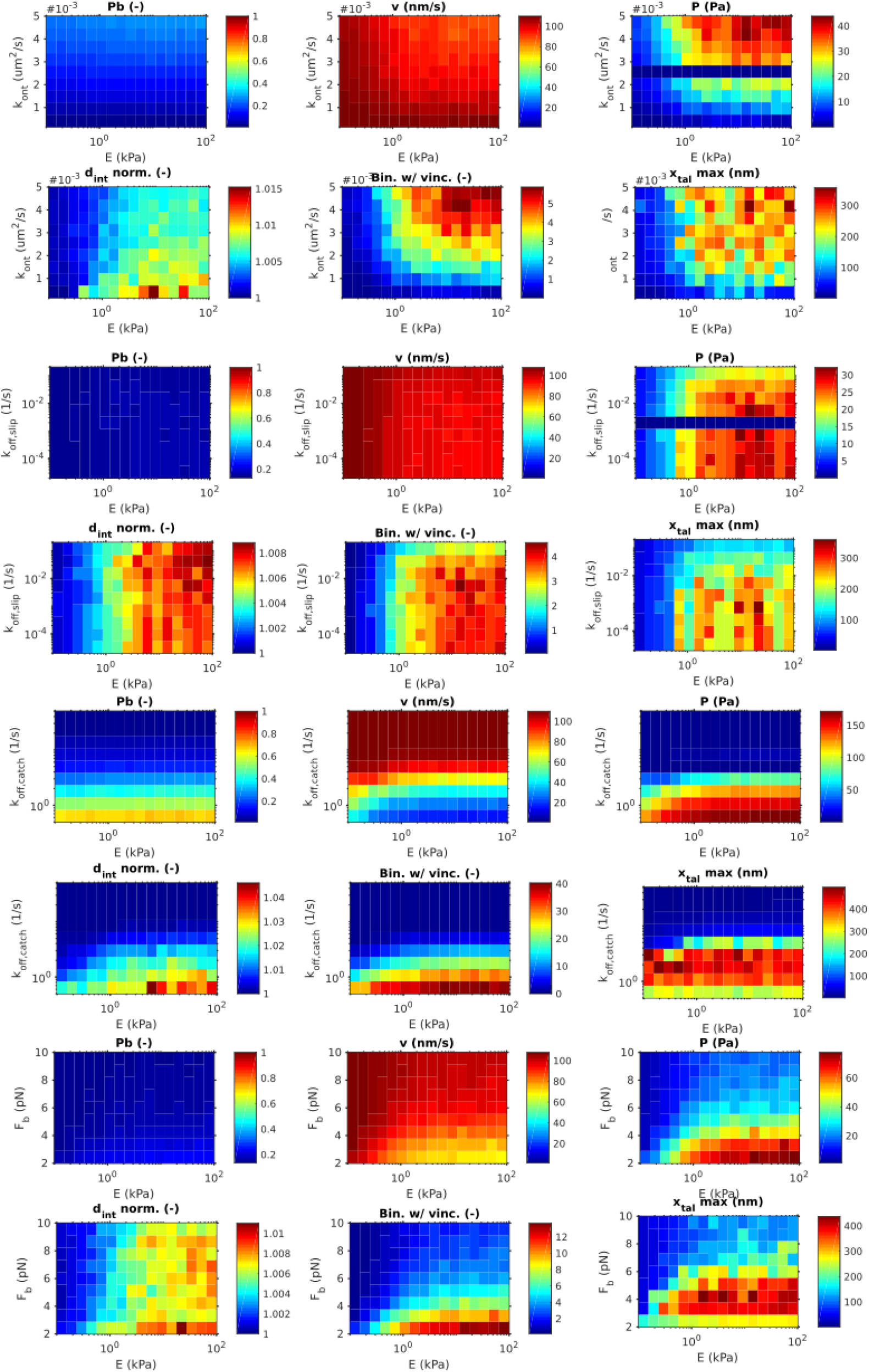
Sensitivity analysis of the model parameters. Sensitivity analysis for the model parameters *k_ont_, k_off,slip_, k_off,catch_* and *F_b,slip_* = *F_b,catch_*. We plot the variables *P_b_, v, P*, *d_int,norm_*, number of binders with vinculin and 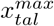 against the Young’s modulus of the substrate *E*. The results are obtained averaging over 10 Gillespie simulations.

To further analyze the effect of the integrins behavior in ACs, we perform a variation of *Fb* and *k_off,catch_* versus *k_ont_* fixing the substrate stiffness to E=10kPa. Because variations of *k_off,slip_* have a small effect in the AC (Fig. 11), we leave it out of the analysis. The results are in agreement with the discussion above (see Fig. A6 for details).

### Variations in AC behavior due to ligand spacing

Finally, we investigate the effect of ligands spacing in the behavior of ACs. As we discussed in the introduction, cell adhesion depends not only on the rigidity of the ECM but also on the spacing between attaching locations. We compute the ligand spacing as a function of the number of ligands in the substrate and of the AC radius (see Fig. A10 for details). We focus on ACs crowded with *α*_5_*β*_3_ integrins [50] and use experimental data of the adhesion length as a function of the substrate stiffness [50] for ligands placed 50, 100 and 200 nm apart. These data and the resulting number of ligands are summarized in Fig. 12. We analyze again the evolution of ACs for stiffness of the substrate in 0.1 — 100 kPa. The remaining parameters are the same as in Section.

**Fig 12.**
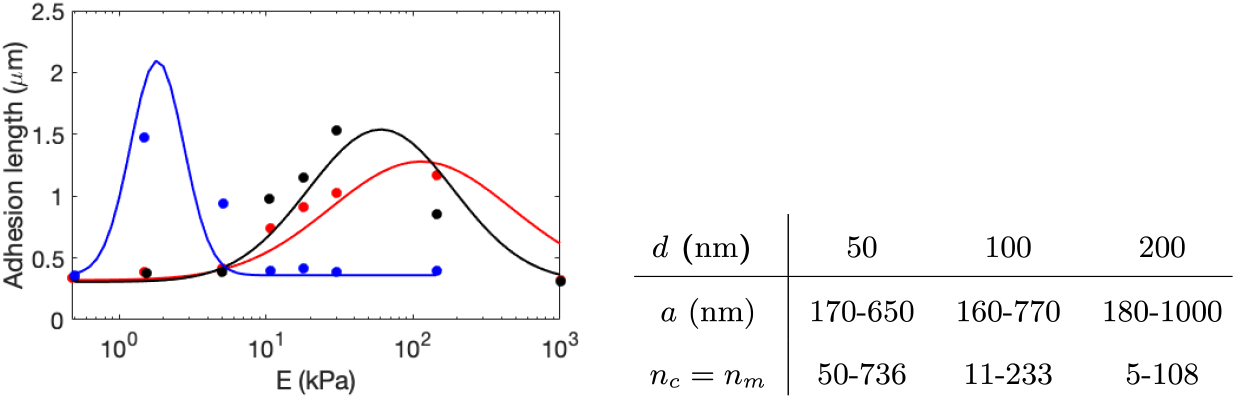
Left: Size of the ACs [50] and fit of the data to a gaussian hill for ligand spacing of 50 nm (red), 100 nm (black) and 200 nm (blue). Right: Number of ligands *n_c_* and radius of the adhesion *a* for the variation of ligand spacing in *α*_5_*β*_1_ integrins.

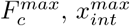 and 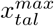 increase as the stiffness of the substrate increases, with slightly higher values for substrates with ligand spacing of 50 nm than of 100 nm (Fig. 13). However, when the ligands are placed 200 nm apart, there is a sharp increase of these variables between 0.5 kPa and 2 kPa, overpassing the values for the 50 nm case at 2 kPa. Between 2 kPa and 10 kPa, we obtain a sudden decrease of the adhesion size down to 180 nm, and just 5 ligands, where it keeps constant. The overall behavior of the bound probability, *P_b_*, the actin velocity, *v*, the average force *F_c,bound_*, the displacements of the integrins, *x_int,bound_*, and talin, *x_talin,bound_*, and the normalized density of integrins *d_int,norm_* is similar for the three distances. These results suggest that, on average, the behavior of the CAMs does not depend on the ligand spacing, although the maximum force and displacements of integrins and talins show striking differences depending on ligands spacing.

The substrate displacement depends on the displacement at each binder and the total number of bound binders. As we obtain a similar average force for the three cases, and a similar probability of bound binders, we obtain larger substrate displacements as we increase the number of ligands (Fig. 13). Likewise, the total number of bound vinculins depends on the number of bound binders as well as on the number of unfolded domains per talin. We show an increase of bound vinculins for the 50 nm and 100 nm cases as the substrate stiffness increases. The bound probability is comparable in these two cases and the stretch of the talin rod, and consequently the unfolding events, is higher in the 50 nm case. This explains the increase up to 80 and 20 bound vinculins for the 50 nm and 100 nm cases, respectively. However, the 200 nm case does not present an increase in bound vinculins in stiff substrates, because of the very low number of bound binders in substrates stiffer than 1 kPa. As previously discussed for ACs crowded with *α*_5_*β*_1_ integrins, the normalized integrin density is not affected by the increase in vinculin activity. Therefore, the effective integrin recruitment for the cases analyzed is mostly negligible and the differences shown in the talin mechanosensing (number of vinculin recruited) have no effect in the traction forces.

Finally, the cell traction increases for increasing ligands number and substrate rigidities. The maximum tractions, found for the stiffest substrates are ≈ 75, 150 and 600 Pa for the 200 nm, 100 nm and 50 nm cases, respectively. Because the cell traction is computed as the ratio of all forces acting on the substrate over the area of the AC, and *P_b_* is similar for the three distances, traction in the ACs increases for increasing number of ligands, which occurs for decreasing distance in a same adhesion area (see values at E = 5 kPa, Fig. 13) or, to a lesser degree, for increasing adhesion areas for constant ligands spacing (Fig. 13). These results indicate that the cell traction at a single AC is mostly determined by the behavior of the single adhesion chains and that the density of bound binders is the determining factor for the cell traction.

**Fig 13.**
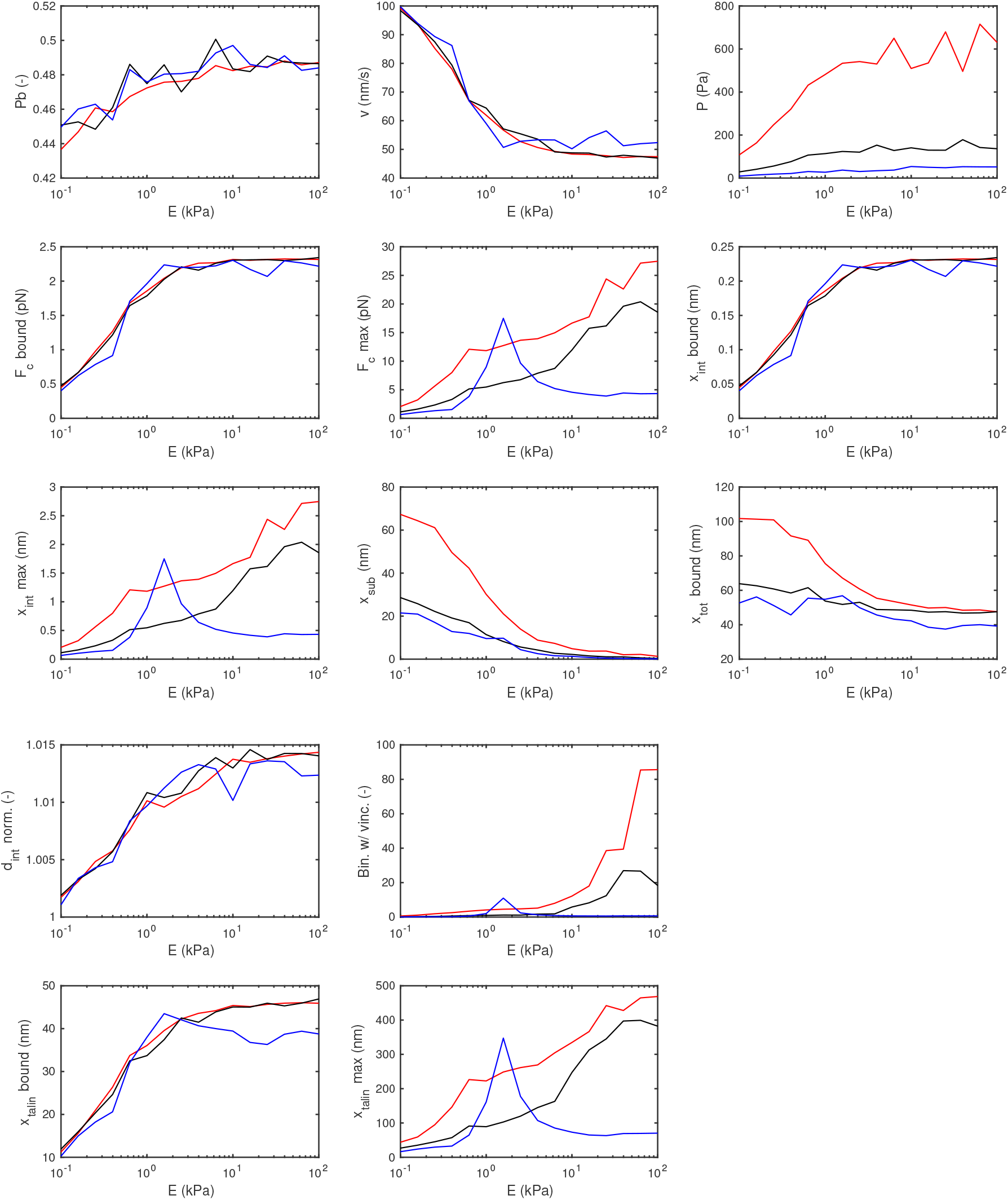
Results for variation of ligands spacing for *α*_5_*β*_1_ integrins. *d* 50 nm (in red), *d* = 100 nm (in black) and *d* = 200 nm (in blue).

## Conclusions

Integrin-based cell adhesion is a central mechanical function to the cell [1]. These cell adhesions are responsible for the attachment of cells to the ECM [3] and, therefore, for controlling cell motility in development, e.g. during neural formation [94], nuclear mechanotransduction [95] and diseases processes such as tumor invasion [96].

In this work, we have first carefully analyzed previous clutch models for cell-ECM dynamics. As previously shown [43, 8, 71], they closely reproduce the cell response at the whole cell-scale. We also identified a number of simplifications and inconsistencies with the mechanical response of previous experimental observations at the scale of the CAMs, which arise from an over-simplified description of the AC. First, the chain of CAMs was reduced to a unique linear spring that roughly gathers the responses of each individual CAMs. This is not just a structural simplification but, more importantly, it results in unphysiological values of the forces and displacements obtained at the molecular chains. Second, the substrate has been traditionally simplified to a single linear spring element that represent the deformation of the entire AC. The weakest link in the adhesion chain is identified in the integrin-fibronectin bond [8, 71]. It is clear that a group of integrins link to several attachment points in the ECM and, therefore, the deformation exerted on the ECM should change pointwise.

To address these issues, we adopted a detailed model of talin [24] that includes the non-linear mechanical behavior and the unfolding and refolding events of each talin domain. Then, we included detailed data on the integrin behavior [40, 69] and the subsequent vinculin binding to the talin rod. Moreover, we modeled the ECM deformation at each binding location by means of Green’s functions. We then integrated all these elements and solved the adhesion dynamics at the whole AC as well as the individual CAM scale.

We used our multi-scale model to understand how ACs crowded with either *α_V_β*_3_ or *α*5*β*1 behave mechanically at both the AC and the molecular levels. Our results agree with previous clutch models and experimental data on traction forces. More importantly, our results on the deformation and force in each CAM closely agree with previous data. This is a key aspect of our model, because it allows us to analyze the whole AC dynamics, while significantly increasing the model predictions at the molecular scale. In other words, we have built the model from the basic components of the AC and, still, the model reproduces the results at the whole adhesion scale. For those parameters that may change depending on specific adhesion cases, such as the substrate stiffness or the type of integrin with which a specific cell is crowded, we ran a sensitivity analysis. This analysis allowed us to specifically pinpoint threshold values at which the traction force would increase or vanish. This is an important aspect in the cell function, because changes in cell traction are associated with alterations in the cell function, including tissue cohesiveness or cell migration.

Our model also showed that ligand distance may not be, by itself, responsible for changes in CAMs response. However, we did show important differences in cell tractions. These differences are due to changes in the AC size. However, previous data on cell tractions for *d* = 50 nm and *d* = 100 nm in human breast myoepithelial cells showed a similar response [50]. The discrepancy between the model results and the experimental data may be due to the spatial scales analyzed. Indeed, we have mostly compared our computational results with experimental data at the whole cell scale (see, e.g., [8, 50, 43]). However, our model specifically tackles single ACs while the experimental traction forces are usually obtained as an averaged quantity over cell sections larger than the actual AC size. The density of ACs per area, and not only the size of the complex, may explain these differences. For example, a FA density higher for *d* =100 nm than for *d* = 50nm would give overlapping curves for the cell traction. We obtained very similar curves for the actin velocity when changing the distance between the ligands, and this is coherent with the experimental results [50]. However, our results refer to the velocity of the actin filaments within a stress fiber, while the experiments measure the retrograde flow velocity. Therefore, experimental data on single ACs will be helpful to validate the model and advance toward a better analysis of the AC behavior. Moreover, data on vinculin binding, integrin density and AC size while varying the substrate stiffness will help us to improve the model predictions.

We believe that we have advanced in the modeling and understanding of single ACs. However, there are still modeling aspects to improve for a better understanding of biological functions of cell adhesion. For example, epithelial cells follow a catch bond behavior with talin reinforcement [8, 71]. Neurons, on the other hand, follow a pure slip behavior [43]. How specifically the type of integrins within the AC impacts the adhesion response across multiple cell types is still poorly understood. To reproduce the traction forces on ACs of cells crowded with *α*5*β*1, we had to increase integrins lifetime 2-fold with respect to the actual values [40]. This inconsistency could be the result of a poor integrin dynamics modeling or missing mechanotransductive aspects in the model. The binding rates of vinculin to talin, *k_onv_* = 1 × 10^8^ s^-1^ was previously adjusted to fit experimental results [8, 71]. Previous data showed a vinculin binding rate of 0.2s^-1^ [97]. Probably, the model does not capture correctly some particular aspect of the AC behavior, which should be addressed, together with further experimental studies.

Our model predicts that force itself cannot be responsible for AC disassembling because, at steady state, the rate of binding and unbinding reach a situation of semi-equilibrum in most of the cases. AC disassembling may be a downstream mechanotransductive result that we did not include in the model. Future studies should also focus on how forces induce mechanotransduction and biochemical processes to control AC disengagement. In line with this, we have considered that the AC remains always of the same size or, if we change it, it is directly imposed in the model based on previous experimental data. The analysis of the change in size during the entire formation and disengagement of the AC should also be addressed in the future.

In summary, we developed a clutch model to analyze single ACs at multiple scales. We have advanced in the understanding on how single CAMs that forms an AC behave. Overall, our model is a step forward in the efforts of rationalizing cell adhesion mechanics and it may also allow us to engineer the cell adhesion response to design better biomimetic tissues [98, 99, 100, 101, 102].

## Methods

### Description of the clutch model

First, the actin flow is simply computed by a force-velocity relation as:

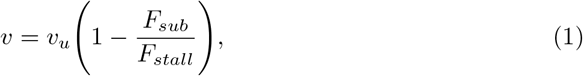

where *v_u_* is the unloaded velocity of the actin flow. The retrograde flow velocity reduces if the ratio between the reaction force in the substrate *F_sub_* and the stall force of the myosin motors, *F_sub_*, increases. In the case of mature ACs, thick stress fibers crowded by myosin motors pull on the adhesion patch with a total stall force *F_stall_* = *n_m_F_m_*, where *F_m_* is the force required to stall the activity of one myosin motor and *n_m_* the number of motors in the system.

Then, the displacement of the engaged clutches is computed as Δ*x* = *v*Δ*t*, where Δ*t* is the time step in the Monte Carlo or Gillespie simulation. The force at each *i* — *th* clutch, *F_c,i_*, is given by

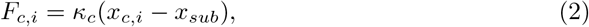

where *κ_c_* is the stiffness of each molecular clutch and *x_c_,i* is the displacement of the *i* – *th* molecular clutch. The substrate is represented by a simple Hookean spring such that the force on the substrate is *F_sub_* = *κ_sub_x_sub_*. The stiffness of the linear spring is *κ_sub_* = *E*4*πα*/9, where *E* is the Young modulus of the substrate and *a* the radius of the AC. Then, the displacement of the substrate is obtained by solving the balance of forces between the *n_eng_* engaged molecular clutches and the substrate as:

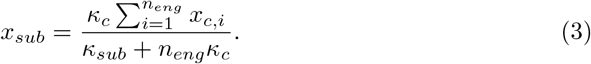

Once *x_sub_* is computed, we can obtain *F_sub_* and the cell traction as *P* = *F_sub_*/*πa*^2^, where (*πα*^2^) is the area occupied by a circular AC.

Once the force at each clutch, *F_c,i_*, is obtained, we compute the binding and unbinding events at each molecular clutch. The *n_c_* molecular clutches are allowed to associate with the ECM, with rate *k_on_*, or to disengage, according to a dissociation rate 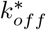. The Bell’s model is the simplest approach for a force-dependent unbinding rate that increases exponentially as

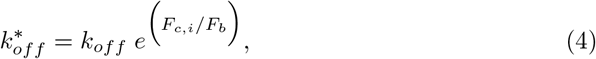

where *k_off_* is the unloaded dissociation rate and *F_b_* is the characteristic bond rupture force. This law follows a slip-behavior, meaning that as the force increases, the lifetime of the bond decreases exponentially (see Fig. 14). Experimental data also showed bonds that follow a catch behavior [40], in which the dissociation rate first decreases and then increases exponentially with the applied force as

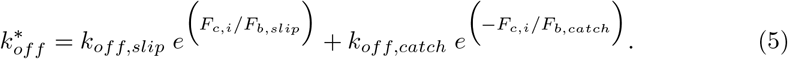

*k_off,slip_* and *k_off,catch_* are the unloaded dissociation rates, and *F_b,slip_* and *F_b,catch_* are the characteristic bond rupture forces, via the slip and catch pathways, respectively. The new bound and unbound clutches are then updated for the next time step when all the relations above are again computed.

However, the catch bond model in Eq. 5 does not take into consideration the the Cyclic Mechanical Reinforcement (CMR) [69]. The CMR increases the lifetime of integrins when they are subjected to cycles of engagement and disengagement before a stable AC is formed. To take into account the effect of CMR, previous models increased the off rate at low forces (< 1 pN) (see Fig. 14) [8] and adopted an unbinding rate with CMR 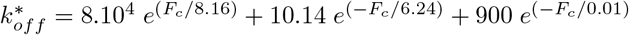. We will use this particular form of 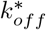 in this section.

**Fig 14.**
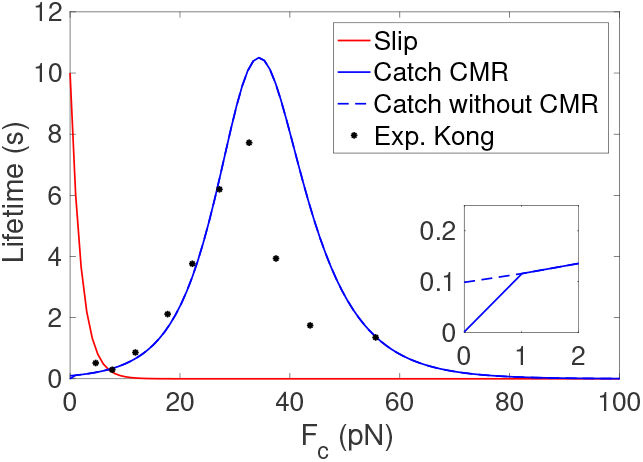
Lifetime of the weakest link for slip (red) and catch bonds with CMR (solid blue). Experimental data of integrin *α*_5_*β*_1_ are shown in dots (black) [69]. The inset shows a zoom of the lifetime for small forces: catch case with CMR (solid blue) and without CMR (dashed blue).

For all simulations in this section, we fix the radius of the AC to *a* =1700 nm. The number of ligands is *n_c_* =1200 in the catch case [8], which results in a ligands spacing of *d* = 100 nm (see Fig. A10 for details). To seemingly compare slip and catch cases, we keep *d* = 100 nm constant for the slip case, and fixing *n_c_* =75 [43] we obtain an AC radius of *a* = 447 nm. The remaining model parameters are summarized in Table 2. We analyze the model results in terms of the substrate stiffness, with Young’s modulus in the range of 0.1 - 100 kPa. This is the stiffness range found in biological tissues [70] and on which most in vitro studies have focused (see, e.g., [43, 8]). We analyze the probability of bound binders *P_b_*, the actin velocity v, the cell traction P, the maximum and the average force over the bound binders, 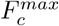 and *F_c,bound_*, the maximum and the average displacement for bound binders 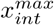 and *x_int,bound_*, and the displacement of the substrate *x_sub_*. We average in time all these variables of interest to obtain the model predictions at specific stiffnesses of the substrate.

### Computation of the clutch model

We use MC simulations with constant time step, Δ*t* =0.005 s. During the simulation time, many events of engagement and disengagement and, often, cycles of formation and rupture of ACs, take place. To evaluate the choice of the final time, we run MC simulations with different final times and choose *t_f_* = 100 s as large enough to achieve accurate results when averaging over time the MC simulations, while reducing at the same time the computational cost of the simulations (see Fig. A3 for details).

The actomyosin network pulls on the adhesion molecules thanks to the contractile forces of the myosin motors. An actin filament is bound to a number of myosin motors, *n_m_*, that generate contractile forces in the molecular chain. The velocity of the actin filament is again *v* = *v_u_*(1 – *F_c_/F_stall_*). *F_c_* is the force at the molecular chain. The stall force is *F_stall_* = *n_m_F_m_*, where *n_m_* = 40 and *F_m_* = 2*pN*, giving a total stall force of 80*pN* per molecular chain (see Appendix,, for details). The total displacement of the engaged adhesion chain is given by *x_tot_* = *v*Δ*t*.

### A multi-scale clutch model

To represent the talin behavior, we incorporate a full-length model of the talin rod mechanics [24], where all the 13 domains have their specific unfolding and refolding rates and a non-linear force-displacement relationship (Fig. 3). We assume that vinculin binds to the unfolded R3 domain, which induces integrins recruitment in the AC, as we described in the previous section. We also assume that actin binds to talin in the ending R13-DD part of the rod. The unfolding rates of the talin rod domains follow a Bell’s model and the folding rates follow an Arrhenius law [84, 24]. Mechanically, the folded domains behave as a freely-jointed chain (FJC), where the end-to-end distance of the folded domain, *x_fol_*, is

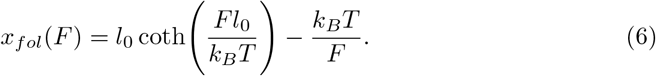

*l*_0_ is the rigid body size of the folded domain and *k_B_T* is the thermal energy. The unfolded domains behave as a Worm-Like Chain (WLC),

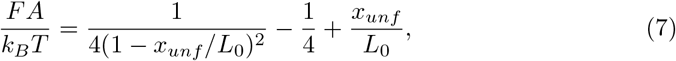

where *x_unf_* is the end-to-end length of the unfolded domain, *A* is the persistence length and *L*_0_ is the contour length. All talin model parameters are described elsewhere [24].

To model the integrin behavior, we use the binding and unbinding rates defined in Section for catch bonds. We use a linear spring to model the force-displacement relation in each integrin molecule, such that *F* = *k_c_*(*x_c_* — *x_sub_*). *x_int_* = *x_c_* — *x_sub_* is the displacement of the integrin molecule and *xc* is the displacement at the integrin-talin bond position.

To mechanically model the extracellular space, we use the Green’s functions in a semi-infinite and isotropic medium [85]. The Green’s function solution relates the elastic displacement in the i-direction at a point x due to a force in the j-direction applied at the origin as

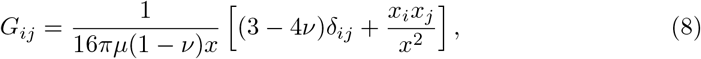

where *μ* is the shear modulus, *v* the Poisson’s ratio of the medium and *x* is the distance from the field point and the point of force application.

We consider a 2D substrate on the XY-plane, where are engagement points have coordinates *x* = (*x, y*). The angle *θ* = 15° between the adhesion chains and the substrate [5, 12] gives forces in the same direction and results in a substrate displacement with non-zero components in the three dimensions. Here, we consider horizontal forces and substrate displacements laying in the XY-plane. Moreover, we consider that all bound binders bear forces with the same direction as all actin filaments in the stress fiber and molecular chains bundle in parallel. Therefore our substrate displacements will have only one dimension and all chains pull on the ECM in the same direction. Therefore, the displacement of the substrate in the direction *i* due to the force also in the direction *i* is given as *x_sub,i_* = *ĜF_c,i_*, where *Ĝ* = *Ĝ_ii_* is the Green’s function referring to the only relevant direction *i*.

We call ***x***_*sub*_ the vector of one-dimensional displacements *x*_*sub*(*s*)_ in all the points *s* inside the FA, and we call ***F***_*c*_ the vector of forces *F*_*c*(*b*)_ in the ligand points *b* = 1,…, *n_c_*. Therefore, the displacement of the substrate at all sampling locations is given as ***x***_*sub*_ = *Ĝ* · ***F***_*c*_.

Once all model variables are solved, the traction exerted by the AC on the substrate is simple computed as *P* = *F/A*. As the Green’s function is singular at the point where the binder is bound and the force is applied, the displacement is computed at a distance of 5 nm from the binder.

### Computation of the multi-scale clutch model

We use here a Gillespie algorithm [86, 87] with variable time step, which is determined by the events rates: binding/unbinding of integrin molecules to the ECM and folding/unfolding rates of the talin rod. The time at which each event, *i*, happens is computed as *t*_i_ = – ln*ξ_i_/k_i_*, for all binders *i* = 1…*n_c_*. *ξ_i_* are independent random numbers uniformly distributed over [0, 1] and *k_i_* is the event rate. After computing the time of all possible events at the current time step, we choose the minimum time *t_i_* and update the corresponding event.

### Number of myosin motors and stall force

In order to find values for the number of myosin motors *n_m_* attached to an actin fibre and the stall force for a single myosin motor *F_m_*, we look for an experimental value of the total stall force *F_stall_* = *n_m_ F_m_*. The ensemble stall force can be written as [103]:

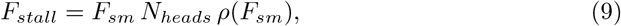

where *F_sm_* = *K_x-bridge_d_step_* is the stall force for a single motor, *N_heads_* is the number of myosin motors and *ρ*(*F_sm_*) is the duty ratio of a single motor at stall, which can be computed as:

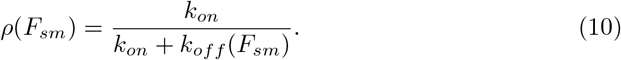

The expression for the myosin off-rate is [104]:

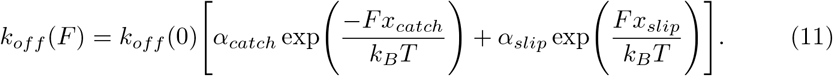

Myosins are a superfamily of motor proteins. Myosin II is responsible for producing muscle contraction in muscle cells in most animals, but it is also found in non-muscle cells inside stress fibres. Hence, it can be further classified into skeletal muscle myosin II, smooth muscle myosin II, and nonmuscle myosin II (NM II) [103]. More precisely, the myosins having a role in cell adhesion are NM IIA and NM IIB. NM IIA mediates the initial maturation of FAs, and NM IIB is found in fibrillar adhesions. Adhesions with NM IIA are dynamic, while those with NM IIB are very stable [105]. In the leading edge of the cell, both types are present, while NM IIA characterizes the rear edge and NM IIB is found in the actin close to the nucleus of the cell [106]. It has been demonstrated that 90% of the traction force generated by mouse embryonic fibroblasts (MEFs) on a fibronectin-coated substrate is lost with the removal of NM IIs, and that NM IIA is responsible for ≈ 60% of the force, whereas NM IIB accounts for the ≈ 30% [107, 108].

Using Eqs. 9, 10 and 11, with parameters in Table 3, we compute the ensemble stall force *F_stall_* for NM IIA and NM IIB. Considering that in our setting there is a percentage of myosin motors of each type, we choose a total stall force in the range 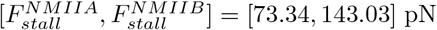, and precisely *F_stall_* = 80 pN. The stall force for a single myosin motor is found to be at least *F_m_* = 1.7 pN [89], therefore, following *F_stall_* = *n_m_ F_m_*, we consider *n_m_* = 40 myosin motors, each with a stall force *F_m_* = 2pN.

**Table 3.** Parameters for myosin motors used inside Eqs. 9, 10 and 11, with experimental references.

| Parameters |  |  |
| --- | --- | --- |
| $K_{x-bridge}$ (pN/nm) | 0.7 | [109] |
| $d_{step}$ (nm) | 5.5 | [109] |
| $N_{heads}$ | 50 | [110] |
| $k_{on}$ (s<sup>-1</sup>) | 0.2 | [111], [112] |
| $k_{off}(0)$ NM IIA (s<sup>-1</sup>) | 1.71 | [111] |
| $k_{off}(0)$ NM IIB (s<sup>-1</sup>) | 0.35 | [112] |
| $\alpha_{catch}$ | 0.92 | [104] |
| $\alpha_{slip}$ | 0.08 | [104] |
| $x_{catch}$ (nm) | 2.5 | [104] |
| $x_{slip}$ (nm) | 0.4 | [104] |

## Acknowledgements

We acknowledge funding from the Spanish Ministry of Science and Innovation (PID2019-110949GB-I00 P.S.), the European Commission (H2020-FETPROACT-01-2016-731957 to C.V. and P.S), the Generalitat de Catalunya (2017-SGR-1278 to P.S. and FI AGAUR 2018 to C.V.).

## Supporting information

**Table A1.** Model parameters and values for the clutch model. For a better comparison, the default values of the model parameters not related to integrin dynamics are the same in both slip and catch cases [8]. For the slip and catch cases, default values and ranges for the parameters used in Figures A7, A8. In order to reduce the number of parameters in the analysis, we neglect the effect of the CMR in the catch bonds (see Fig. 14). Therefore, we do variations over the four model parameters of the catch bond model in Eq. 5 and over the two parameters in the slip bond model in Eq. 4. In terms of the sensitivity range, we vary the number of ligands *n_c_* in two different ways while keeping the radius of the adhesion fixed (i.e., changing the ligand spacing): keeping *n_m_* fixed to the default value, and setting *n_c_* = *n_m_* in order to have one myosin motor available per ligand. For the stiffness of the molecular clutch *k_c_*, we only vary its value downwards because *k_c_* = 1000 pN/nm is already an upper bound of a real value. We take a narrow range of values for *v_u_*. As the stall force is the product of *F_m_* and *n_m_*, we vary the stall force of a single myosin motor *F_m_* while keeping constant the number of myosin motors *nm*. However, *F_m_* has been measured to be ≈ 2 pN so we only make small variations around that value. We take a range of values for the binding rate *k_ont_* considering the data from previous experimental work [8]. Linear or exponential distribution are adopted inside the ranges. The exponential distribution is chosen for ranges covering many orders of magnitude, in order to get more values in the first half of the range.

| Parameters | Slip |  |  | Catch |  |  |
| --- | --- | --- | --- | --- | --- | --- |
|  | Default | Min | Max | Default [8] | Min | Max |
| $E$ (kPa) | - | 0.1 | 100 | - | 0.1 | 100 |
| $a$ (nm) | 1700 | - | - | 1700 | - | - |
| $n_c$ | 1200 | 50 | 2320 | 1200 | 50 | 2320 |
| $\kappa_c$ (pN/nm) | 1000 | 0.01 | 1000 | 1000 | 0.001 | 1000 |
| $k_{ont}$ ($\mu\text{m}^2/\text{s}$) | $2.11 \times 10^{-4}$ | $3.5 \times 10^{-6}$ | $8.3 \times 10^{-4}$ | $2.11 \times 10^{-4}$ | $3.5 \times 10^{-6}$ | $8.3 \times 10^{-4}$ |
| $d_{int}$ (int / $\mu\text{m}^2$) | 300 | - | - | 300 | - | - |
| $k_{off,slip}$ ($\text{s}^{-1}$) | 0.1 | $1 \times 10^{-5}$ | 0.01 | $7.9938 \times 10^{-4}$ | $1 \times 10^{-5}$ | 0.01 |
| $F_{b,slip}$ (pN) | 12 | 0.1 | 20 | 8.1633 | 0.1 | 20 |
| $k_{off,catch}$ ($\text{s}^{-1}$) | - | - | - | 10.143 | 0.1 | 30 |
| $F_{b,catch}$ (pN) | - | - | - | 6.2461 | 0.1 | 20 |
| $F_m$ (pN) | 2 | 0.1 | 5 | 2 | 0.1 | 5 |
| $n_m$ | 800 | 50 | 2320 | 800 | 50 | 2320 |
| $v_u$ (nm/s) | 110 | 20 | 250 | 110 | 20 | 250 |

## Supplementary figures

All the figures of the model results in time for all substrates rigidities described in Section for *α*_5_β_1_ integrins are uploaded in here.

All the figures of the model results in time for all substrates rigidities described in Section for *α*_5_*β*_3_ integrins are uploaded in here.

All the figures of the model results in Section are uploaded in here.

All the figures of the model results in Section are uploaded in here.

**Table A2.** Model parameters of the reinforced case. For the catch model with talin reinforcement, default values and ranges for the parameters used in Fig. A9. We vary the initial density of integrins *d_int_* and the density of integrins added at each recruitment *int_add_* within two orders of magnitude. We do not vary the maximum integrin density, *m_r_*, because it is never reached. We take a range of values to the left of the default value of the vinculin binding rate *k_onv_* [8] because it is several orders of magnitude larger than values reported in literature of ≈ 1s^-1^ [97]. Linear or exponential distribution are adopted inside the ranges.

| Parameters | Catch |  |  |
| --- | --- | --- | --- |
|  | Default<br>[8] | Min | Max |
| $E$ (kPa) | - | 0.1 | 100 |
| $a$ (nm) | 1700 | - | - |
| $n_c$ | 1200 | 50 | 2320 |
| $\kappa_c$ (pN/nm) | 1000 | 0.001 | 1000 |
| $k_{ont}$ ($\mu\text{m}^2/\text{s}$) | $2.11 \times 10^{-4}$ | $3.5 \times 10^{-6}$ | $8.3 \times 10^{-4}$ |
| $d_{int}^0$ (int / $\mu\text{m}^2$) | 300 | 10 | 1000 |
| $k_{off,slip}$ ($\text{s}^{-1}$) | $7.9938 \times 10^{-4}$ | $1 \times 10^{-5}$ | 0.01 |
| $F_{b,slip}$ (pN) | 8.1633 | 0.1 | 20 |
| $k_{off,catch}$ ($\text{s}^{-1}$) | 10.143 | 0.1 | 30 |
| $F_{b,catch}$ (pN) | 6.2461 | 0.1 | 20 |
| $F_m$ (pN) | 2 | 0.1 | 5 |
| $n_m$ | 800 | 50 | 2320 |
| $v_u$ (nm/s) | 110 | 20 | 250 |
| $k_{onv}$ ($\text{s}^{-1}$) | $1 \times 10^8$ | 1 | $1 \times 10^8$ |
| $int_{add}$ (int / $\mu\text{m}^2$) | 24 | 5 | 300 |
| $m_r$ (int / $\mu\text{m}^2$) | 15000 | - | - |

**Fig A1.**
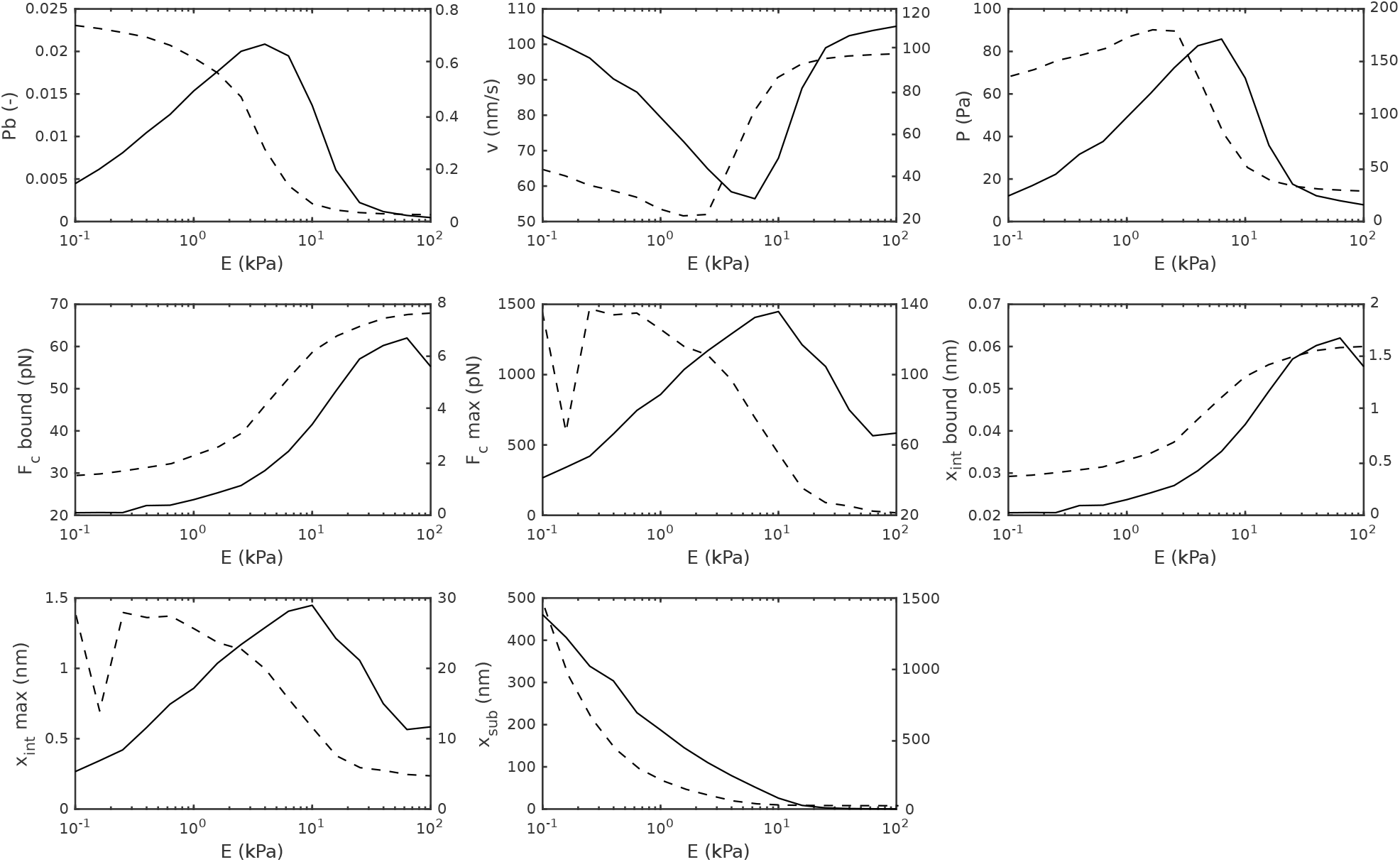
Effect of slip and catch bonds in cell adhesion behavior. Model behavior for slip (dashed line, with corresponding y-axis on the right) and catch (solid line, with corresponding y-axis on the left) cases. Plot against Young’s modulus of the substrate *E* of the variables *P_b_, v, P, F_c_* over bound binders, 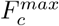, *x_int_* over bound binders, 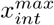 and *x_sub_*. In the slip case, the probability of bound binders, *P_b_*, decreases as the substrate stiffness increases. However, *P_b_* shows a biphasic relation in the catch case, where it first increases and then decreases with an increasing stiffness of the substrate. In absolute values, *P_b_* almost doubles in the catch case with respect to the slip case. At large stiffnesses, the frictional slippage explains the reduction in the number of bound binders. The different *P_b_* at low stiffnesses, where the load-and-fail cycle emerges, is due to the differences in the slip/catch bonds behavior. The maximum lifetime of slip bonds is at low forces (≈0-5 pN), while it maximizes at intermediate forces (≈30-35 pN) for catch bonds. Therefore, if the force at each clutch is low at low substrate rigidities, the model predicts larger binding/unbinding rates for the slip bonds than for catch bonds, which explains the difference in the amount of bound binders at low stiffnesses. The same arguments explain the behavior of the maximum force 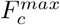 and maximum displacement 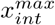 on the molecular clutches. The deformation of the substrate, *x_sub_*, decreases as the stiffness of the substrate increase, and vanishes at high stiffnesses because the increase in the mean force is not enough to compensate the increase in substrate rigidity. The mean force, 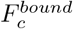, and the mean displacement, 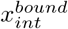, increase as the substrate stiffness increases. However, the maximum and average forces and displacements are one order of magnitude higher for the slip case than the catch case. The hill shapes in *P_b_* and *F_sub_*, as well as their magnitudes, explain the shape of the tractions, where the catch case shows a quick increase in cell traction at low rigidities and lower smaximum values with respect to the slip case. When the cell traction is analyzed against the substrate stiffness, there is an optimal rigidity value that indicates the stiffness at which the cell traction is maximum. The differences in the optimal stiffnesses are controlled by the lifetime of the slip and catch bond dynamics. The retrograde flow *v* follows the same tendency but opposite to the cell tractions. The velocity is maximum when the cell traction is zero (no adhesion) and, therefore, there is no resistance to the flow, while the velocity is minimum when the tractions are maximum (i.e. at the optimal rigidity) and the resistance to the actin flow is maximum. In both slip and catch cases, the optimal rigidity obtained in the clutch model (Fig. A1) is in agreement with previous experimental and theoretical results [43, 8]. Below the optimal rigidity, the load-and-fail cycle rules. As the stiffness of the substrate increases, the average force at the bound binders increases (see below) and so does the traction.

**Fig A2.**
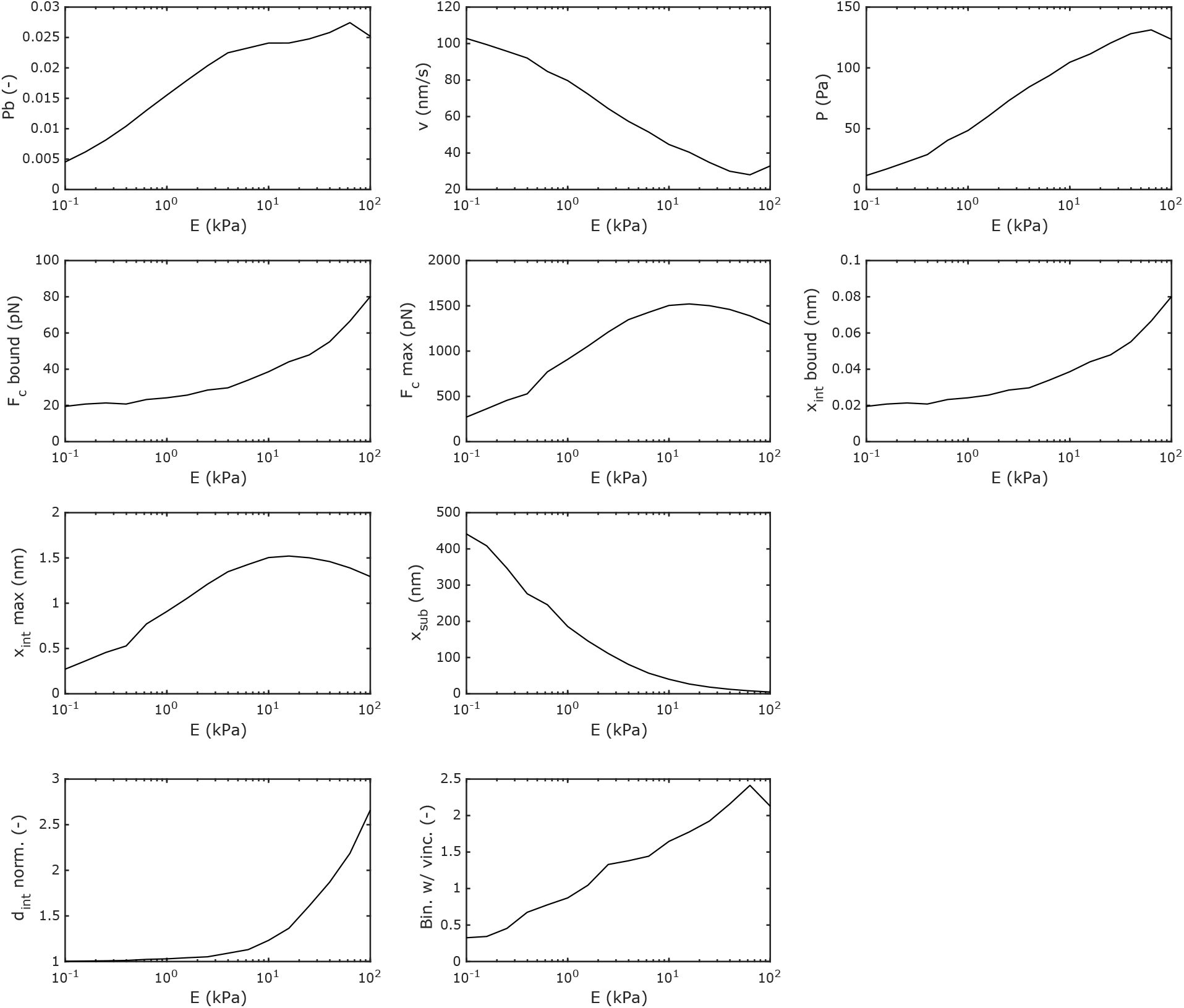
Effect of reinforcement by talin unfolding in cell adhesion. For the model with talin reinforcement, plot against Young’s modulus of the substrate *E* of the variables *P_b_, v, P, F_c_* over bound binders, 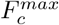, *x_int_* over bound binders, 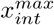, *x_sub_, d_int,norm_* and # binders with vinculin. All variables increase monotonically as the substrate stiffness increases, including the cell tractions, in agreement with previous data, except for the actin velocity *v* and the substrate displacement *x_sub_* instead decrease monotonically. As in previous cases at low substrate stiffnesses, the forces at each molecular chain are low, talin mechanosensing does not activate and, as a result, integrin recruitment does not occur. Therefore, results are comparable to the catch case above A1. In stiff substrates, above, ≈ 10 kPa, we see an increase in all the variables that decreased in the catch case, including the cell tractions. As the substrate stiffnesses increases, the forces at each molecular chain increase, which activates talin unfolding. If talin exposes its VBS, vinculin binds to the talin rod and fosters integrins recruitment within the AC, i.e. the integrin density *d_int_* increases, and so does the binding rate, which eventually increases the probability of bound binders and the cell traction *P*. This process explains the adhesion reinforcement by talin mechanosensing [8, 71].

**Fig A3.**
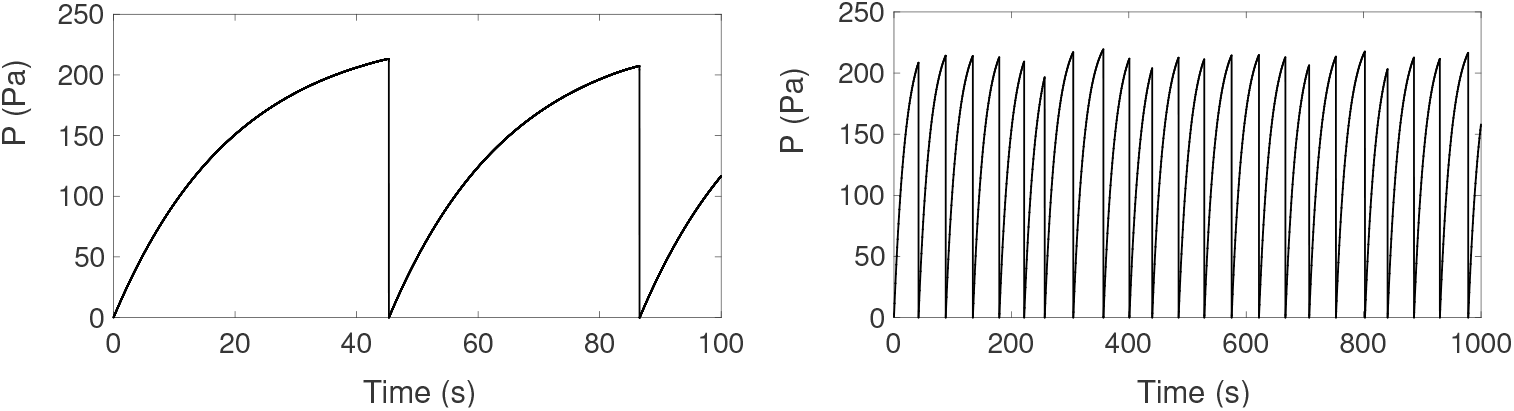
Comparison of the number of MC simulations for two different final times *t_f_* = 100 s and *t_f_* = 1000 s. We run an slip case and fix the Young’s modulus of the substrate *E* = 0.1 kPa. As the MC simulations are concatenated, in order to do the average over MC, we do the average in time of the variable. For *t_f_* = 1000 s, the average of the cell traction is *P* ≈ 138 Pa, while for *t_f_* = 100 s the average is *P* ≈ 132 Pa.

**Fig A4.**
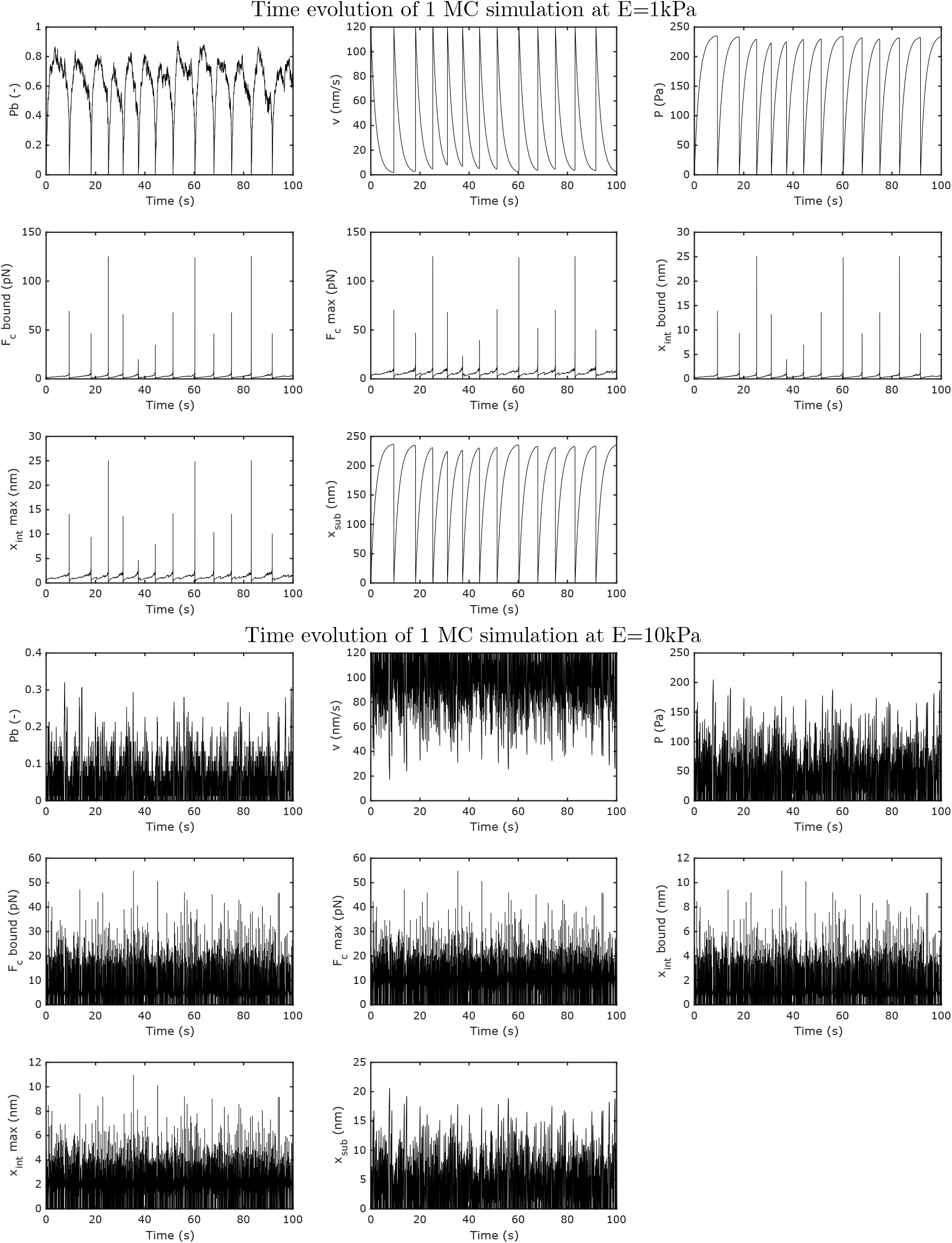
Time evolution of the slip case. Time evolution of the slip case for the variables *P_b_, v, P, F_c_* over bound binders, 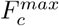, *x_int_* over bound binders, 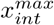 and *x_sub_*, for *E* = 1kPa and *E* = 10 kPa. Model parameters are given in Table 2. The lifetime of the adhesion clusters are considerably shorter for stiffer substrates.

**Fig A5.**
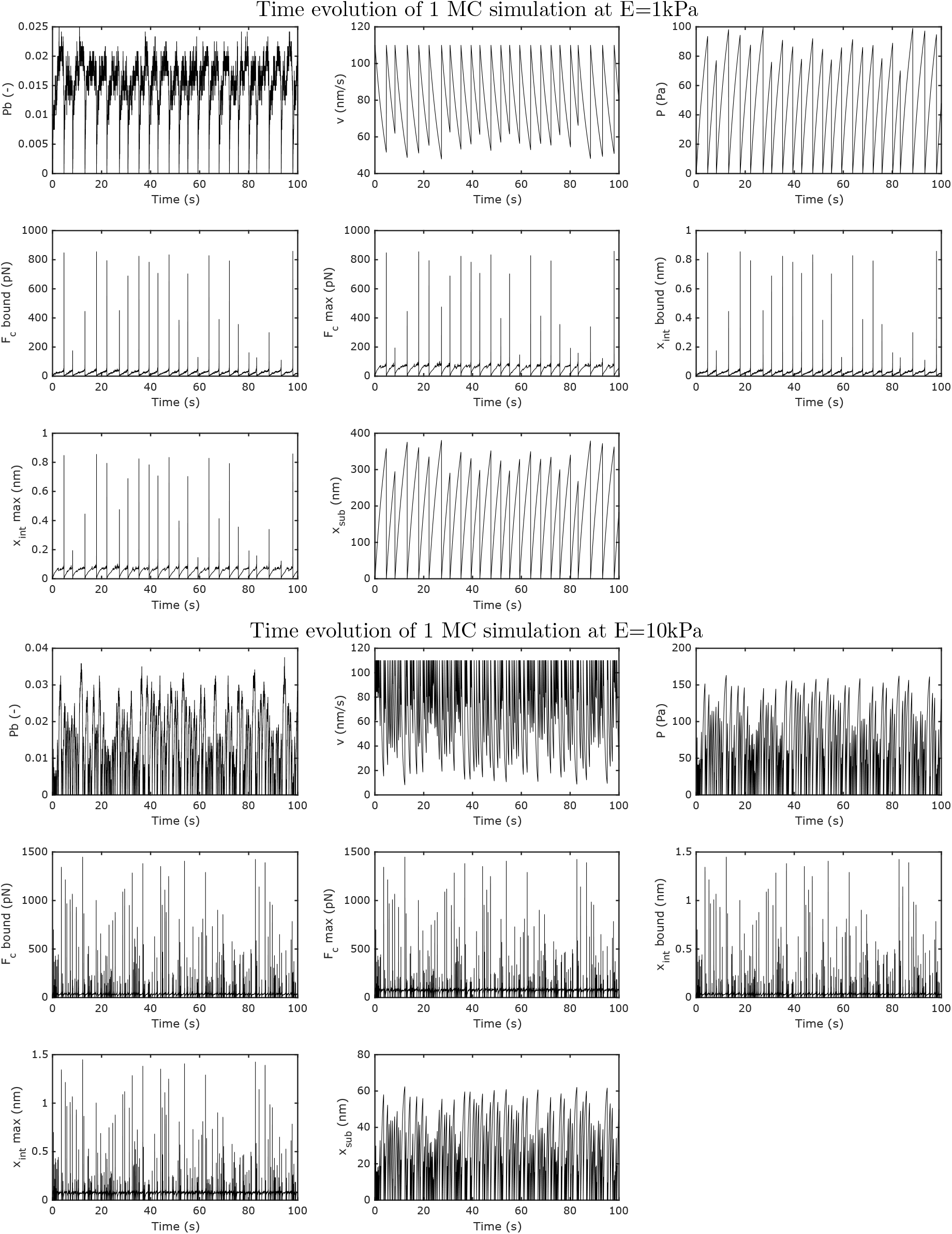
Time evolution of the catch case. Time evolution of the catch case for the variables *P_b_, v, P, F_c_* over bound binders, 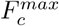, *x_int_* over bound binders, 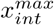 and *x_sub_*, for *E* =1 kPa and *E* = 10 kPa. Parameters from Table 2. The MC loops to get a cluster completely made of free binders are considerably shorter for stiffer substrates.

**Fig A6.**
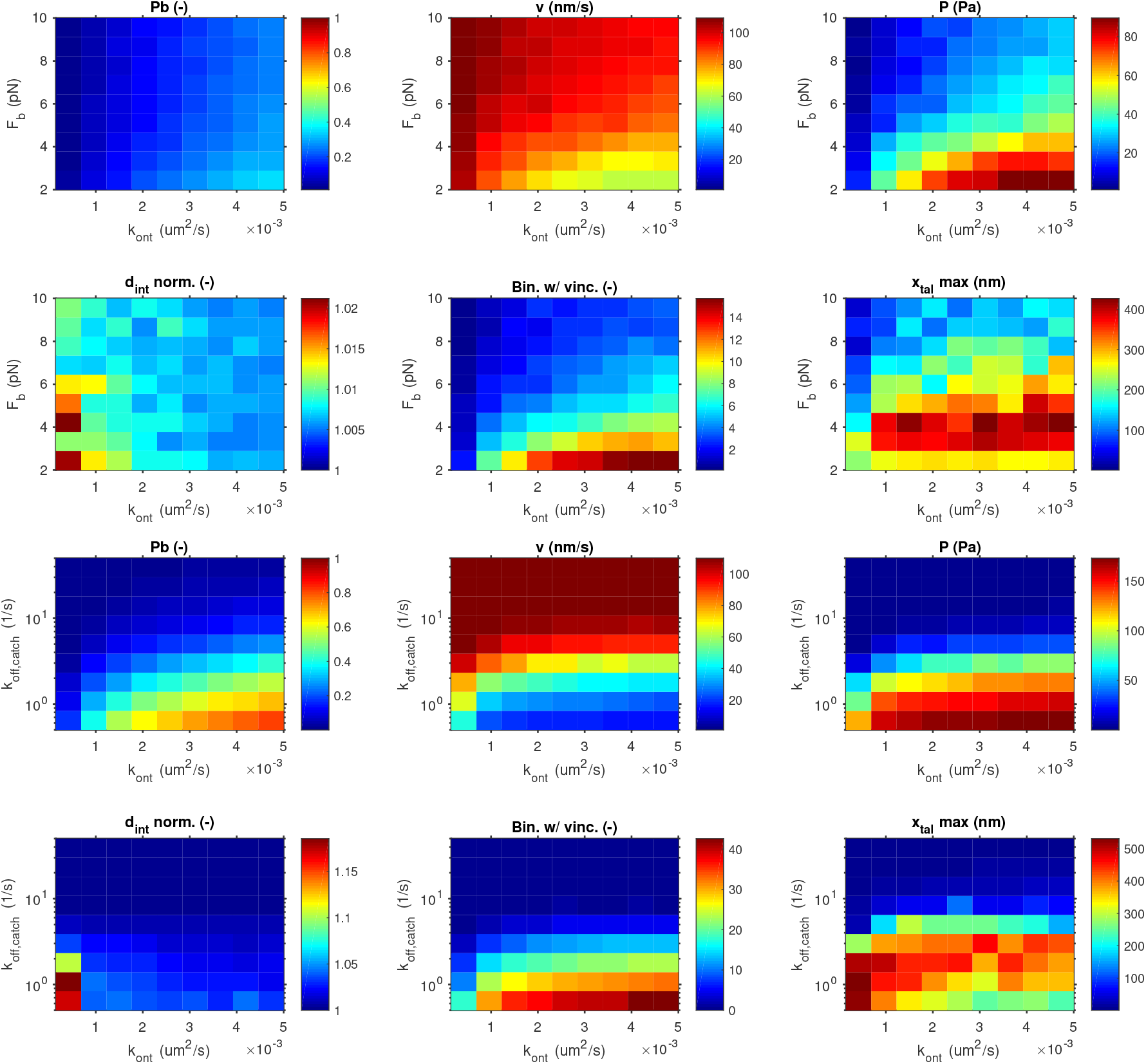
Sensitivity analysis of the clutch model. Fixing the Young’s modulus of the substrate E=10 kPa, plot of the main variables varying simultaneously the parameters *k_ont_* and *F_b,catch_* = *F_b,slip_*, and the parameters *k_ont_* and *k_off,catch_* later. The combination of the highest *k_ont_* and the smallest *k_off_* results in the maximum number of bound binders, bound vinculins and, effectively, in cell traction. It is also worth to note that the maximum tractions are not obtained for the maximum talin displacement. A similar result can be found for the combination of the highest *k_ont_* and the smallest *F_b_*, the characteristic bond rupture force through both the slip and catch pathways. Indeed, considering negligible the slip pathway because of the orders of magnitude in the range of *k_off,slip_* compared to those of *k_off,catch_*, a variation in the force *F_b_* has the same impact as a variation in *k_off,catch_*. Similarly to the observations for the standard clutch model, an increase in *k_off,catch_* induces a fast unbinding of all bound binders even at low forces, which results in low tractions, low mechanosensation of the talin rod and recruitment of vinculing and an increase of the retrograde flow. On the other hand, an increase in the *k_ont_* value results in the opposite effects, creating more traction, reducing the retrograde flow and increasing the unfolding dynamics of the talin rod. The increase of the rupture force, *F_b_*, has a similar impact to that observed by the increased *k_off,catch_*. Overall, these results are in agreement with previous results.

**Fig A7.**
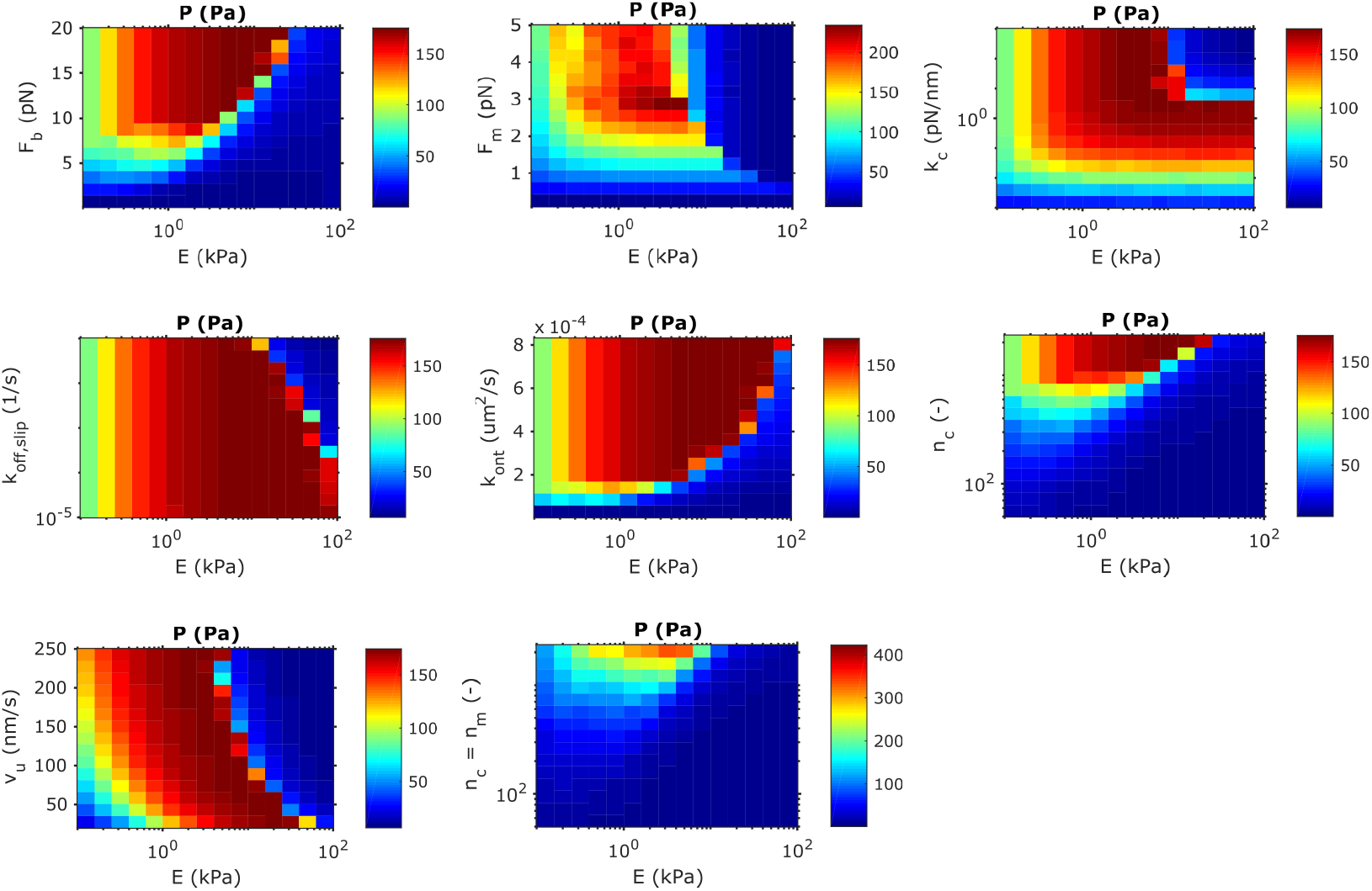
Sensitivity analysis of the slip case of the clutch model. Results of the cell traction *P* against Young’s modulus *E* and varying in a suitable range the parameters *F_b_, F_m_, κ_c_*, *k_off,slip_*, *k_ont_*, *n_c_*, *v_u_*, *n_c_* kept equal to *n_m_*, *k_off,catch_*, *F_catch_*. Default values and range is shown in Table A1. An increase in *k_off,siip_* makes the binders of the AC to break faster for increasing applied forces, which decreases the cell traction. In the slip case, there is no effect in cell tractions at low substrate stiffnesses, below the optimum stiffness, because the force is weakly transmitted to the clutches. The cell traction *P* corresponding to the optimal stiffness, located at ≈7-10 kPa, has the same value for all choices of *k_off,slip_*. The effect of the increase of *k_off,slip_* can only be observed above the optimal stiffness, where the forces mostly deforms the clutches. As *F_b,slip_* controls the characteristic bond rupture force, a decrease in *F_b,slip_* decreases the off-rate. Therefore, the cell traction *P* corresponding to the optimal stiffness increases as *F_b,slip_* increases. When *k_ont_* increases, the cell traction *P* at the optimal stiffness increases. This is expected as an increase in *k_ont_* increases the amount of bound binders in the AC. In terms of the clutch rigidity, we observed an increase in the traction force up to ≈1*pN/nm* as we increase *k_c_*, while its value remains constant in E. An increase in *F_m_* keeps the optimum stiffness in ≈2-3 kPa and that the maximum traction is obtained for *F_m_* = 2 —3pN, while it drastically vanishes below that value because the motors do not have enough force to deform the system. An increase in *n_c_* logically produces an increase in the cell traction *P* at the optimal stiffness, which slightly shifts toward higher stiffnesses, because of more bound binders. Similarly, by increasing *n_c_* = *n_m_*, however, we observe an increase in the cell traction *P* at the optimal stiffness, which shifts toward higher stiffness, although less evident than in the case in which *n_c_* is varied independently of *n_m_*. Here, as we increase the number of binders, we also increase the number of myosin motors, increasing the pulling forces. Finally, an increase of *v_u_* does not change the value of *P* at the optimal stiffness but shifts it to lower rigidities because, when the flow velocity increases, the force transmission rises, and the optimal rigidity shifts toward lower values where the decrease in substrate rigidity can accommodate the increasing force rates.

**Fig A8.**
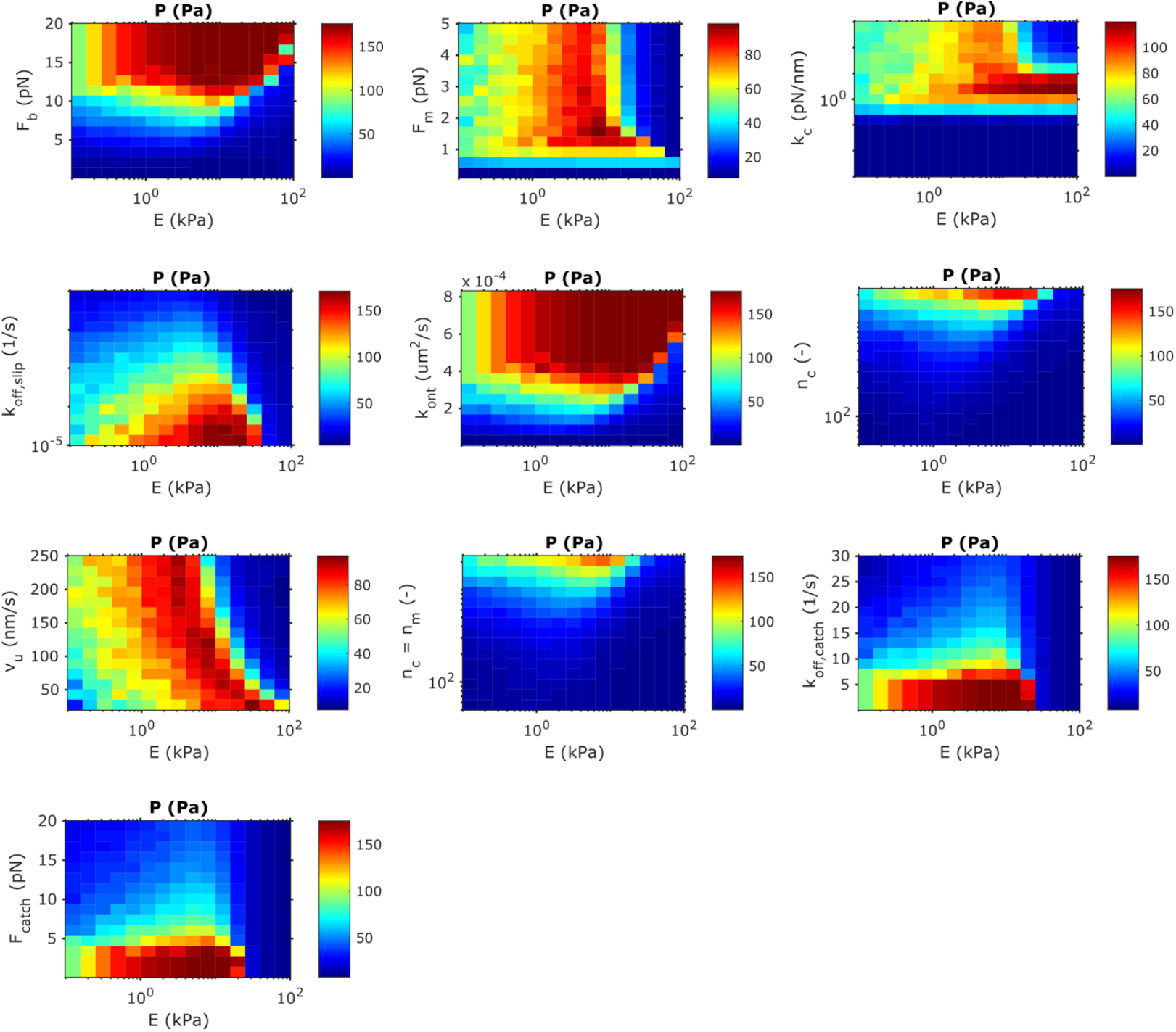
Sensitivity analysis for the catch case of the clutch model. Results of the cell traction *P* against Young’s modulus *E* and varying in a suitable range the parameters *F_b_, F_m_, κ_c_, k_off,slip_, k_ont_, n_c_, v_u_, n_c_* kept equal to *n_m_, k_off,catch_, F_catch_*. Default values and range is shown in Table A1. The cell traction *P* corresponding to the optimal stiffness decreases for increasing values of *k_off,slip_* until a total disengagement of ACs, and the optimal stiffness is slightly shifted toward soft substrates. At low substrate stiffnesses, i.e. low force transmission, the catch behavior dominates, and the complex is mostly disengaged. An increase in *k_off,catch_* reduces the catch effect in the bond and, consequently, the lifetime of bound binders. Indeed, we see a 4-fold decrease in the cell traction *P* at the constant optimal stiffness of ≈10 kPa. As *F_b,slip_* controls the characteristic bond rupture force, a decrease in *F_b,slip_* decreases the off-rate. Therefore, the cell traction *P* corresponding to the optimal stiffness increases as *F_b,slip_* increases. The opposite behavior is obtained for increasing values of *k_off,catch_*. When *k_ont_* increases, the cell traction *P* at the optimal stiffness increases. This is expected as an increase in *kont* increases the amount of bound binders in the AC. In terms of the clutch rigidity, we observed an increase in the traction force up to *≈1pN/nm* as we increase *k_c_*, while its value remains constant in E. The traction force is very low (≈1 *pN/nm*). At ≈1 *pN/nm,* the traction force increases monotically up to its maximum values at the maximum substrate stiffness analyzed. Above ≈1(pN/nm), we see a different response: the optimal stiffness shifts to ≈10kPa, from where tractions quickly drop to zero. The response of the traction as we vary *F_m_, n_c_* and *v_u_* are comparable to the results in the slip case.

**Fig A9.**
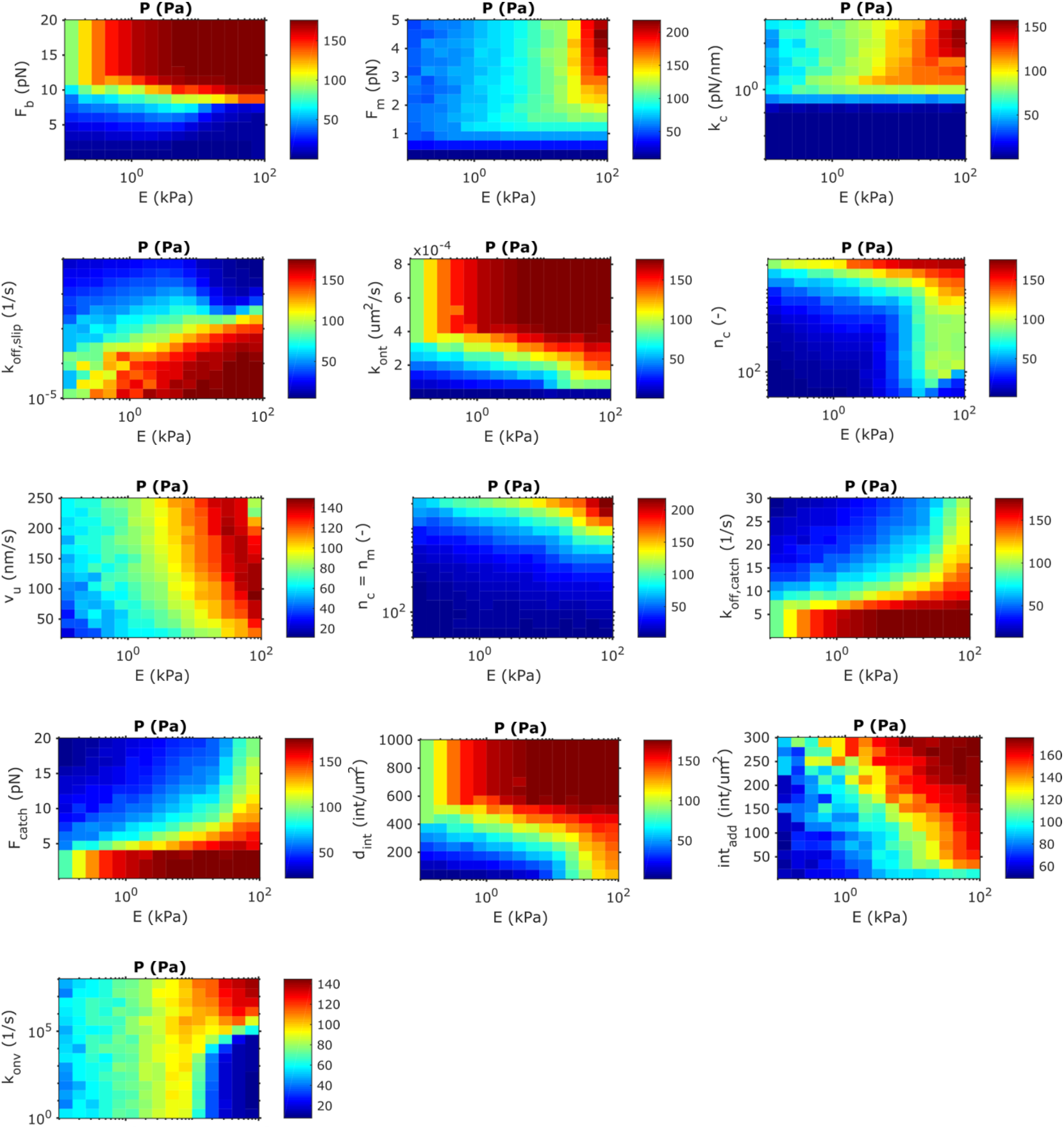
Sensitivity analysis of the reinforced case. For the model with talin reinforcement, values of the cell traction *P* following colormap, against Young’s modulus *E*, varying in a suitable range the parameters Fb, *F_m_*, *κ*_c_, *k_off,slip_, k_ont_, n_c_, V_u_, n_c_* kept equal to *n_m_, k_off,catch_, F_catch_*, 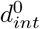, *int_add_, k_onv_*, as in Table A2. The results show a response to the model parameters similar to the non-reinforced model for low stiffnesses of the substrate, where the talin reinforcement has not taken place due to the low force in each clutch (see the main text). For the upper part of the ranges and all model parameters, the cell traction *P* always increases as the Young’s modulus of the substrate increases. This change with respect to the drop in traction forces for the catch case is due to the integrin recruitment when vinculin binds to the unfolded talin. In other words, the system never gets into the frictional slippage regime. In terms of the parameters that control the integrin recruitment, we see that increasing *int_add_* the cell traction *P* corresponding to the optimal stiffness increases. This is because the integrin density added at each recruitment step increases the total integrin density and, consequently, the on-rate of integrin binding increases. Similarly, an increase in the on-rate of vinculin to talin, *k_onv_*, results in an increase in the cell traction *P* in the upper part of the stiffness range, while lower stiffnesses there is no effect. This is due to the fact that integrin recruitment is only activated once talin is unfolded, which only occurs at high stiffnesses. We also see that integrin recruitment is only effective for values of *k_onv_* > 10^5^. Below this value, integrin recruitment does not happen and we observe a decrease in cell traction similar to the catch case without reinforcement (see main text).

**Fig A10.**
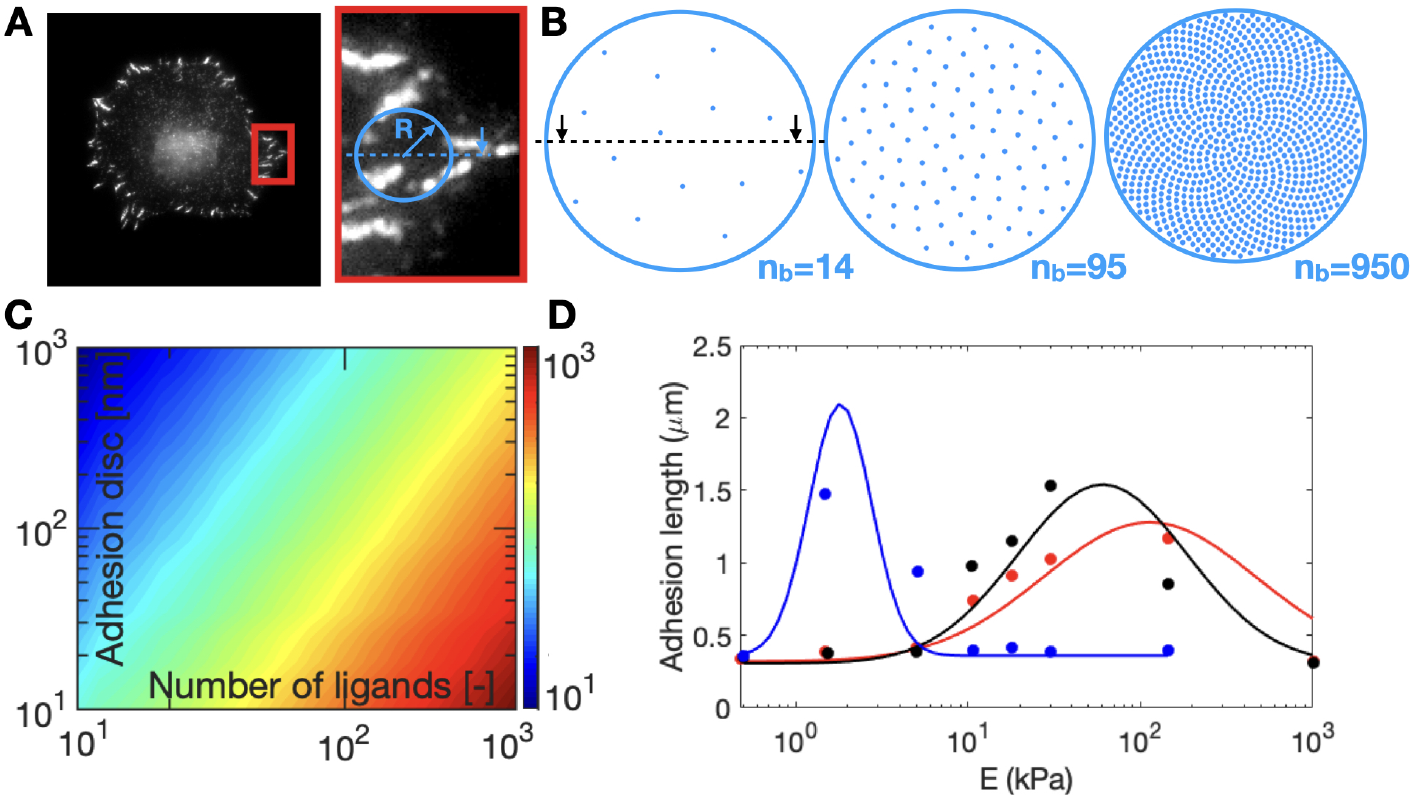
Number of equispaced ligands *n_c_* as a function of the adhesion radius. We compute a set of *n* distributions for *n_c_*=1 to *n_c_*=5000 for a radius of *μm.* Then we extend the set by scaling each one of the distributions by different number of radii, m, from 10nm to 1μm. Altogether, we obtain a set of nxm distributions of *n_c_* ligands within adhesion of varying radii. (a) Cell expressing AC. Inset of the front cell. (b) Different number of ligands in an AC of the same area. (c) Computation of the ligands distance (color map) as a function of the adhesion size and number of ligands. Among all the combinations of number of ligands and radii, we obtain sets of ligand distances that are not physical viable. Integrins have an approximate equivalent radius of 2.5nm, based on an estimated footprint of the integrin dimer [9]. Considering a dense packing of the integrins, the maximum density is ≈ 25,000 integrins/*μm*^2^. Therefore, integrins can not be apart less than ≈ 5nm, which we impose as a physical constraint in the spacing between ligands. Furthermore, we also constrain the spacing by the number of actin filaments that fit in a given domain of radius R. Assuming a stress fiber up to 30 actin filaments, an average filament diameter of 20 nm and a distribution almost parallel to the contact plane, there is a limit of bound ligands distance of 20 nm. In ventral-like adhesions, the angle of actin and talin with respect to the membrane has been measured in 2-6 and 15°, respectively [5]. In order to accommodate the actin filaments within the stress fiber and assuming a stoichiometry of talin-integrin of 1:1 and talin to actin of 1:1 to 1:3 (three talin domains are able to link to actin filaments) we get a range of possible ligand distances that goes up to ≈ 400 nm. (d) Experimental data [50] and fitting of the adhesion size for different ligands spacing as a function of the substrate stiffness.

## Notes

### Competing Interest Statement

The authors have declared no competing interest.

